# Developmental alterations of indirect-pathway medium spiny neurons in mouse models of Huntington’s disease

**DOI:** 10.1101/2024.05.10.593545

**Authors:** Margaux Lebouc, Léa Bonamy, Jakob Scharnholz, Quentin Richard, Gilles Courtand, Jérôme Baufreton, Maurice Garret

**Author notes:** these authors share seniority on the study (Jérôme Baufreton, Maurice Garret).

## Abstract

Huntington’s disease (HD) is an inherited neurodegenerative disorder caused by a mutation in the gene encoding the Huntingtin protein (Htt). While symptoms, primarily characterized by progressive deterioration of the striatum and motor and cognitive functions, typically manifest in adulthood, recent studies have also highlighted developmental defects in HD. Indeed, alterations in cortical and striatal development have been observed in individuals carrying the mutation as early as in embryonic stages. However, despite the striatum being one of the most affected regions in HD, few studies have investigated potential developmental alterations in this structure, especially in the early weeks after birth. To address this question, we compared striatal development between wild-type (WT) mice and two murine models of HD, R6/1 and CAG140 mice crossed with reporter mice to identify D1- and D2-expressing medium spiny neurons (D1- and D2-MSNs). Using *ex vivo* electrophysiology and neuronal reconstruction, we observed that the maturation of electrical properties was selectively disrupted in D2-MSNs of the matrix compartment of HD mice during the first post-natal days. D2-MSNs arbor also an increased dendritic complexity. When studying the establishment of striatal afferents, we observed that cortico-striatal glutamatergic transmission was specifically reduced in D2-MSNs during the second postnatal week. All these alterations were transient before the circuit normalized on its own after the second postnatal week. These anatomical and electrophysiological data highlight the significant impact of the Htt mutation on numerous striatal development processes during the postnatal period. Interestingly, we observed that these alterations specifically affect MSNs in the indirect pathway. This preferential vulnerability aligns with the early death of these neurons in adulthood, suggesting that early treatment of these alterations could potentially modify the disease’s progression.

## Introduction

Huntington’s disease (HD) is caused by a dominant mutation in the gene encoding the huntingtin protein (Htt), which leads to degeneration in striatal and cortical structures. The symptoms of HD, characterized by motor, cognitive, and behavioral changes, typically have a delayed onset, occurring around the age of 45 (Ross et al., 2019; Tabrizi et al., 2022). However, evidence suggests that HD mutation also affects the early stages of brain development (for review,(Lebouc et al., 2020; Ratié & Humbert, 2024; van der Plas et al., 2019)

Numerous studies have demonstrated the crucial role of Htt protein in development. Initial research showed that a complete deletion of HTT gene is lethal in the embryonic stage (Duyao et al., 1995; Nasir et al., 1995; Zeitlin et al., 1995). HD mouse models have revealed that mutated Htt (mHtt) disrupts neural progenitor cell division and neuronal migration to cortical layers resulting in reduced cortical thickness (Barnat et al., 2017, 2020; Molina-Calavita et al., 2014). Additionally, mHtt impacts the maturation of cortical neurons at the postnatal stage by impairing their dendritic maturation and axonal growth (Barnat et al., 2017; Capizzi et al., 2022; McKinstry et al., 2014). During the first postnatal week, glutamatergic transmission appears diminished in the cortex of the HdhQ111 mouse model coupled with increased excitability of pyramidal neurons. However, these alterations are transient, with circuitry normalizing during the second postnatal week (Braz et al., 2022). Furthermore, either expressing mHtt or reducing normal Htt expression during developmental stages triggers HD-like alterations in adulthood and thus strongly suggests that these developmental changes may contribute to HD pathogenesis (Arteaga-Bracho et al., 2016; Molero et al., 2016).

Recently, evidence has revealed developmental alterations in humans carrying the HD mutation. These findings show a reduction in proliferative cells and premature increases in cortical neuronal progenitors in 13-week-old human fetuses with the HD mutation (Barnat et al., 2020). Moreover, striatal hypertrophy and an alteration of the developmental trajectory of the striatum have been reported in children aged from 6 to 18 years, who are carriers of the mutation. (van der Plas et al., 2019). Additionally, hyperconnectivity between the cerebellum and striatum has been reported in these HD carrier children (Tereshchenko et al., 2020). As all these changes are perceptible from the earliest age tested (6 years), suggesting an even earlier striatal development alteration (Lebouc et al., 2020). Taken together these breakthrough studies in humans validate the neurodevelopmental alteration hypothesis previously suggested by animal model studies.

Despite the well-established histopathological alterations observed in the striatum of human adult HD carriers or mouse models (Pouladi et al., 2013; Vonsattel et al., 1985), research on possible alterations in striatal development is limited. The striatum, the main input structure of the basal ganglia, is divided into patches and matrix compartments as delineated by neurochemical markers (J. B. Smith et al., 2016). Both compartments predominantly contain medium spiny neurons (MSNs) divided into two sub-populations: one expressing the D1 dopamine receptor (D1-MSNs), and the other expressing the D2 receptor (D2-MSNs). These two intermingled compartments and MSN subtypes are involved in different neuronal circuits (Flaherty & Graybiel, 1994; Friedman et al., 2015). In human HD brains, striatal D2-MSNs are affected earlier than D1-MSNs (Albin et al., 1992). But does mHtt affect striatal circuit formation during the early postnatal period? This critical period shapes the morphological and electrophysiological properties of striatal neurons, along with their connections with the rest of the brain. The maturation of neuronal activity and glutamatergic transmission during this time appears essential for maintaining a functional circuit into adulthood (Braz et al., 2022). Thus, disturbances observed in the striatum in adult HD could result from developmental disruptions occurring in the early postnatal days.

Here, we used R6/1 and CAG140 mice crossed with D1- or D2-GFP reporter mice as animal model of HD to assess the development of striatal MSNs during the first two post-natal weeks. Electrophysiological and morphological properties of D1- and D2-MSNs were studied. Afferent connections from the motor cortex were also assessed.

## Results

### The number and the distribution of D1- and D2-MSNs are not altered in R6/1 mice

Homozygous D1- and D2-GFP mice were crossed with R6/1 mice to generate WT and R6/1 littermates. To identify D1- and D2-MSNs, we conducted multiple immuno- labeling assays to assess the expression of GFP, Ctip2, or MOR in brain sections of D1- and D2-GFP mice. Ctip2 serves as a specific nuclear marker for all striatal MSNs, while GFP is expressed specifically in D1-MSNs and D2-MSNs, respectively. MOR was used as striosome marker (Pert et al., 1976; Tinterri et al., 2018).

Analyses on brain sections from WT::D1-GFP at postnatal days 0, 3, 6, 8, 10, and 15 (n=1-3 per genotype and age, **Figure S1A-G**) revealed that the percentage of GFP^+^ neurons among Ctip2^+^ MSNs was initially low in the earliest ages (5 ± 1.5%, 6 ± 0.1%, 11 ± 2.5%, 22 ± 2.3%, 26%, and 42% at P0, P3, P6, P8, P10, and P15, respectively). In R6/1::D1-GFP littermate mice, the percentage was comparable (5 ± 0.3%, 6 ± 0.9%, 13 ± 2.6%, 21 ± 3.7%, 31%, and 46%), showing no significant difference from that of WT littermates at corresponding ages. Between P0 and P6, the GFP labeling was restricted in MOR-rich territories i.e. in the striosome compartment (**Figure S1A-G**).

Similarly, analyses of WT::D2-GFP (n=2 per genotype and age, **Figure S2A-F**) showed that 40 ± 0.8% of Ctip2^+^ MSNs were GFP^+^ at P0, reaching a plateau at P3, P6, P8, and P10, where the percentage of GFP^+^ MSN was 46 ± 0.04%, 46 ± 1%, 45 ± 0.8%, and 44 ± 5.3%, respectively. In R6/1::D2-GFP mice, the percentage was 41 ± 0.8%, 45 ± 0.02%, 44 ± 1.8%, 43 ± 2.8%, and 45 ± 2.4%, with no significant difference from that of WT littermates at the corresponding ages.

In addition, based on quantification of the total number of Ctip2^+^ neurons, we compared the evolution of MSN density between WT and R6/1 mice (**Figure 1A-F**). MSN density decreases progressively with age as striatal volume increases in agreement with the literature showing that during the first postnatal week, 30% of striatal neurons die, drastically reducing neuronal density (Dehorter & Del Pino, 2020; Fishell & van der Kooy, 1991). The MSN density between WT and R6/1 mice, was not significantly different (P>0.05, Two-way ANOVA). Moreover, as the D2-GFP line discriminates D1-MSNs from D2-MSNs from the first postnatal days, we measured the specific density of these two subpopulations (**Figure 1G-H**). In the same way, we observed a progressive decrease in the density of these neurons (WT, D1: P0: 4.88 ± 0.05 x10^3^ neurons per μm², P10: 1.89 ± 0.45 x 10^3^ neurons per μm; D2: P0: 3.30 ± 0.08 x10^3^ neurons per μm², P10: 1.44 ± 0.03 x 10^3^ neurons per μm) without detecting any significant difference between the two genotypes. Thus, these results suggest that the Htt mutation does not impact the overall distribution of D1-MSNs and D2-MSNs within the striatum during the first two postnatal weeks.

**Figure 1:**
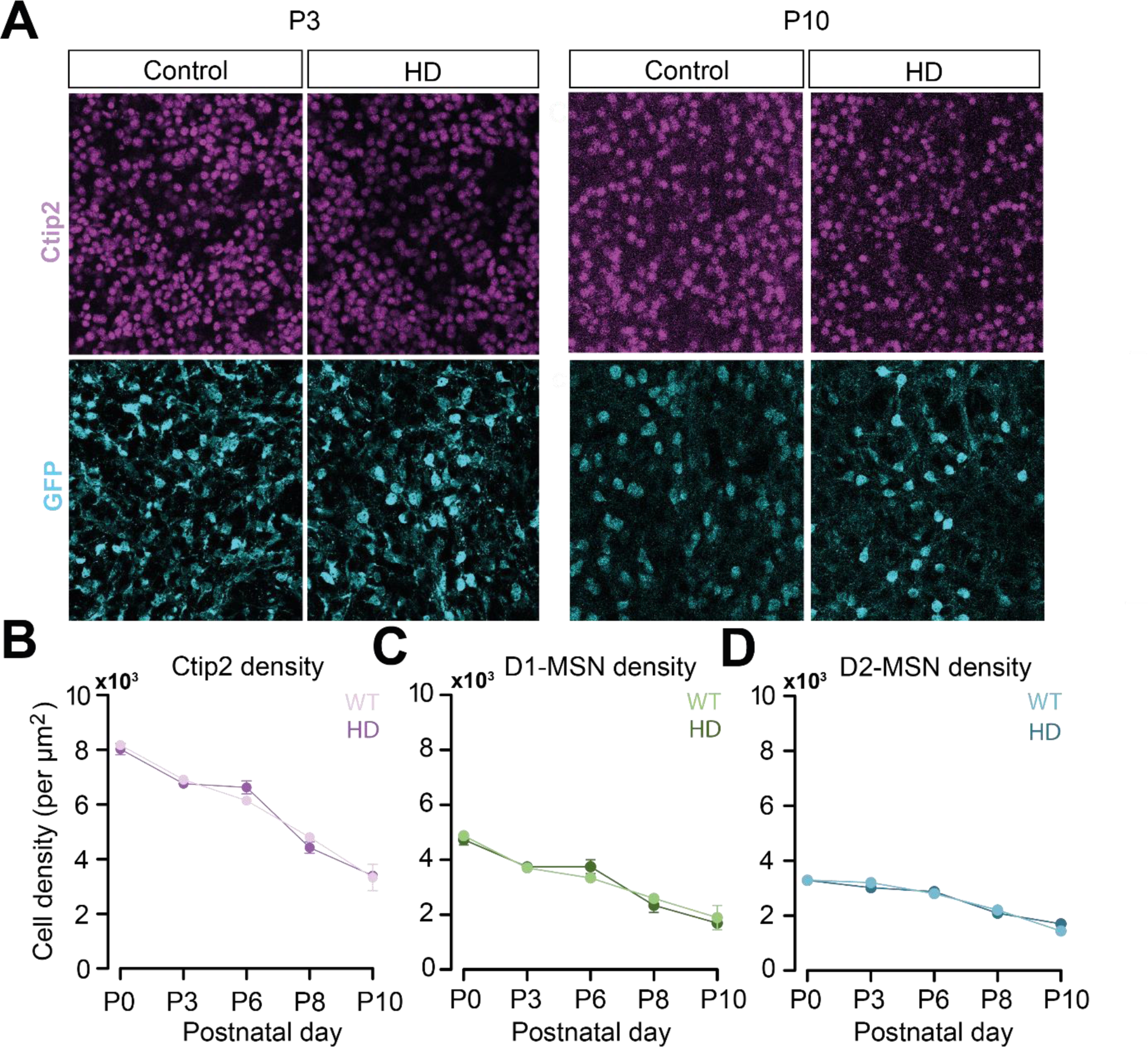
Neuron density is unchanged in R6/1 mice. **A.** Illustration at P3 and P10 of immunohistochemistry against Ctip2 a selective marker of MSNs and GFP which was used to label D2-MSNs in WT::D2-GFP and R6/1::D2-GFP mice at P0, P3, P6, P8, and P10. **B-D**. Density of Ctip2^+^ neurons (**B**), D1-MSNs Ctip2^+^ GFP^-^ (**C**) and D2-MSNs GFP^+^ (**D**) per μm² shown longitudinally.

### The electrophysiological properties of D2-MSNs appear to be altered in HD mice during the early postnatal period

To determine if the Htt mutation impacts the functional properties of the MSNs, we carried out *ex vivo* electrophysiological recordings on coronal slices from WT or R6/1 mice at 4 postnatal stages: P0-2, P3-6, P7-10 and P15-16. The recorded neurons were filled with biocytin for post-hoc identification and reconstruction. To differentiate between recorded D1-MSNs and D2-MSNs in D1-GFP mice we used the transcription factor Ebf1, which is specifically expressed by matrix D1-MSNs (Tinterri et al., 2018) and preproenkephalin (PPE), which is specifically expressed by D2-MSNs (Krajeski et al., 2019; Lee et al., 1997) (**Figure 2A**). These markers show age-dependent variations in expression. Thus, in WT::- and R6/1::D1-GFP in the first postnatal days (P0-6), we employed both Ebf1 and GFP to identify all D1-MSNs. Conversely, neurons that were negative for both GFP and Ebf1 were considered D2-MSNs (**Figure 2A**). However, as Ebf1 expression is mainly associated with embryonic stages and declines rapidly after P5 (Tinterri et al., 2018), from P7 onwards, we relied on the PPE protein to identify D2-MSNs (**Figure 2A**). In parallel, D1-MSNs and D2-MSNs recorded from WT::- and R6/1::D2-GFP mice were identified as GFP-negative (GFP^-^) or GFP-positive (GFP^+^) neurons, respectively. In all cases, we also employed the Ctip2 marker to confirm that all recorded neurons were indeed MSNs and not striatal interneurons. Since data obtained on D1-MSNs from both D1- or D2-GFP mice were not significantly different, they were pooled together. Similarly, we combined the data on D2-MSNs from both types of mice.

**Figure 2:**
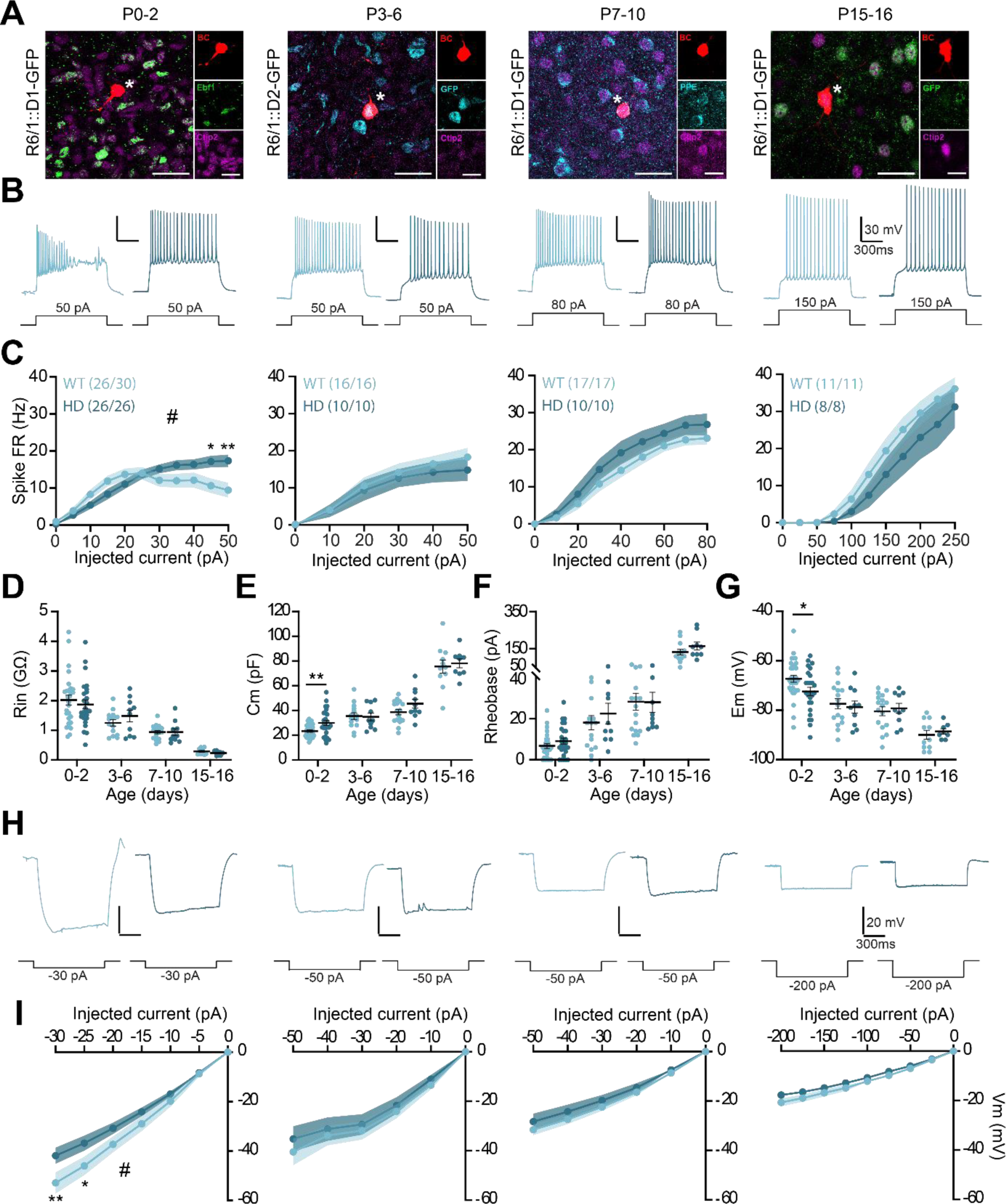
Maturation of the electrophysiological properties of D2-MSNs is impaired in R6/1 mice during the first postnatal days. **A**. Examples of recorded biocytin-filled neurons at 4 postnatal stages: P0-2, P3-6, P7-10 and P15-16. By performing immunohistochemistry against the marker for MSNs, Ctip2, and the specific markers for either D1-MSNs, Ebf1, or D2-MSNs, PPE, we were able to confirm the D2 nature of the recorded neurons. Recorded neurons are identified by an asterisk. **B**. Representative responses of D2-MSNs in WT mice (light blue) or R6/1 mice (dark blue) following injection of a depolarizing current. **C**. Mean discharge frequency/injected current curves obtained for each age. **D-G**. Main electrophysiological properties of D2-MSNs with membrane input resistance (Rin, **D**), capacitance (Cm, E), rheobase (**F**) and resting membrane potential (Em, **G**) plotted longitudinally. **H**. Representative responses of D2-MSNs in WT mice (light blue) or R6/1 mice (dark blue) following injection of a hyperpolarizing current. **I**. Potential/current curves obtained for each age. Scale bars: 25 and 10 μm.

We first assessed neuronal excitability by injecting various depolarizing currents (**Figure 2B**). In WT mice, we observed that most neurons (26/30, 87% of neurons) at the P0-2 developmental stage were capable of initiating low-amplitude action potentials. However, they exhibited immaturity as they were unable to discharge at high frequencies in response to larger currents. Additionally, some neurons (5/30, 17% of neurons) displayed spontaneous activity. As age increased, we noted a higher threshold for inducing a similar discharge frequency, consistent with a progressive increase in rheobase (**Figure 2C**). Furthermore, we observed a progressive maturation of membrane properties in these neurons, including a decrease in membrane resistance (**Figure 2D**), an increase in capacitance (**Figure 2E**), and a hyperpolarization of the resting membrane potential (WT: P0-2: -67.3 ± 1.4 mV, P3-6: -77.438 ± 2.1 mV, -80.5 ± 1.7 mV and P15-16: -90 ± 1.8 mV; **Figure 2F**). These observations align with the expected maturation of electrophysiological properties of D2-MSNs reported in the literature (Dehorter et al., 2011; Krajeski et al., 2019).

In contrast to WT mice, D2-MSNs in R6/1 mice were able to initiate action potentials at P0-2 (26/26, 100% of neurons). Moreover, the evoked discharge frequency in R6/1 mice appeared significantly different from that in WT mice (#, F^(10, 500)^ = 4.651; P<0.0001; repeated measures ANOVA; **Figure 2C**). More specifically, this difference was observed at the highest currents, at 45 and 50 pA (*p<0.05, Sidak’s multiple comparison test). Unlike neurons from WT mice, D2-MSNs from R6/1 mice are still capable of generating high-frequency action potentials in response to high currents. However, the number of spontaneously active cells remains the same (5/26 neurons, 19% of neurons).

Significant differences were also observed in the membrane properties at P0-2. Indeed, D2-MSNs from R6/1 mice showed a significantly increased membrane capacitance compared to WT mice (WT, n=30, Cm = 22.98 ± 1.03 pF; R6/1, n=26, Cm = 30.04 ± 2.07 pF; p=0.0027, Student’s t test; **Figure 2E**). In addition, their resting potential appeared more hyperpolarized than that of WT mice (WT, Em = -67.3 ± 1.4 mV; R6/1, Em = -75.4 ± 1.7 mV; p = 0.0221, Student’s t test; **Figure 2G**). In contrast, membrane resistance and rheobase showed no significant change in R6/1 mice at P0-2 (p>0.05; Student’s t test; **Figure 2D, F**). These alterations are found specifically at P0-2, since from P3 onwards, neither discharge frequency (p>0.05, ANOVA) nor membrane parameters (p>0.05, Student’s t test) differed between the two genotypes.

By injecting different levels of hyperpolarizing current, we observed the gradual establishment of inward rectifier current mediated by Kir channels (**Figure 2H**). MSNs are known to express Kir potassium channels, which are activated by hyperpolarization to maintain a constant membrane potential. These channels are progressively established during the postnatal period (Dehorter et al., 2011). During the first postnatal days, neurons hyperpolarized in proportion to the injected current intensity, resulting in a linear potential/injected current curve (**Figure 2I**). This linearity suggests that Kir channels are not yet present at this developmental stage, thus not influencing neuronal potential. As age progresses, the rectifying current becomes established, as indicated by the progressive change in the curve’s shape, becoming less linear. Consequently, the hyperpolarization induced by current injection becomes weaker (WT, P0-2: For -30 pA injected: -52.9 ± 4.3 mV; P3-6: For -50 pA injected: -40.5 ± 5.4 mV; P7-10: For -50 pA injected: -31.6 ± 2.2 mV; P15-16: For -200 pA injected: -20.8 ± 1.6 mV). We found that the potential/injected current curve of R6/1 mice differed significantly from that of WT mice at P0-2 (#, F^(6, 321)^ = 4.043; p=0.0006; repeated measures ANOVA; **Figure 2I**). Specifically, for the strongest negative currents, at -25 and -30 pA, the hyperpolarization induced by current injection was significantly reduced in D2-MSNs from R6/1 mice (*p<0.05, Sidak’s multiple comparison test). This result suggests the presence of a rectifier current, suggesting accelerated expression of Kir channels in D2-MSNs from R6/1 mice. From P3 onwards, no significant difference was observed between genotypes (**Figure 2I**). In conclusion, we observed abnormal maturation of electrophysiological properties of D2-MSNs from R6/1 mice during the first postnatal days (P0-2). These alterations appear to be transient, disappearing after a few days. We compared the electrophysiological properties of D1-MSNs as we did for D2-MSNs and found no difference (**Figure S3**). We also performed the same electrophysiological characterization of D2-MSNs in the CAG140 mouse model of HD (**Figure S4**), in which we observed almost the same alterations of excitability as described in R6/1 mice, such as an increased excitability (**Figure S4B-C**), and a non-linear response to hyperpolarizing current (**Figure S4H-I**) at P0-2 in CAG140 mice. The input resistance (**Figure S4D**) and the rheobase (**Figure S4E**) were also significantly different between WT and CAG140 animals but with a shift towards pups aged P3-6.

### Alterations in the electrophysiological properties of D2-MSNs appear to be compartment-dependent

To investigate whether the observed alterations in D2-MSNs from R6/1 mice at P0-2 were specific to a striatal compartment, we distinguished between striosomal and matrix neurons. Exploiting the bias of the D1-GFP line, we targeted GFP^-^ neurons surrounded by GFP^+^ neurons for striosomal D2-MSN recordings, confirmed by their localization in a DARPP-32-enriched region. Conversely, we targeted GFP^-^ neurons outside the GFP^+^ neuron area for matrix D2-MSN recordings, identified using Ebf1 and DARPP-32 markers (**Figure 3A, B**).

**Figure 3:**
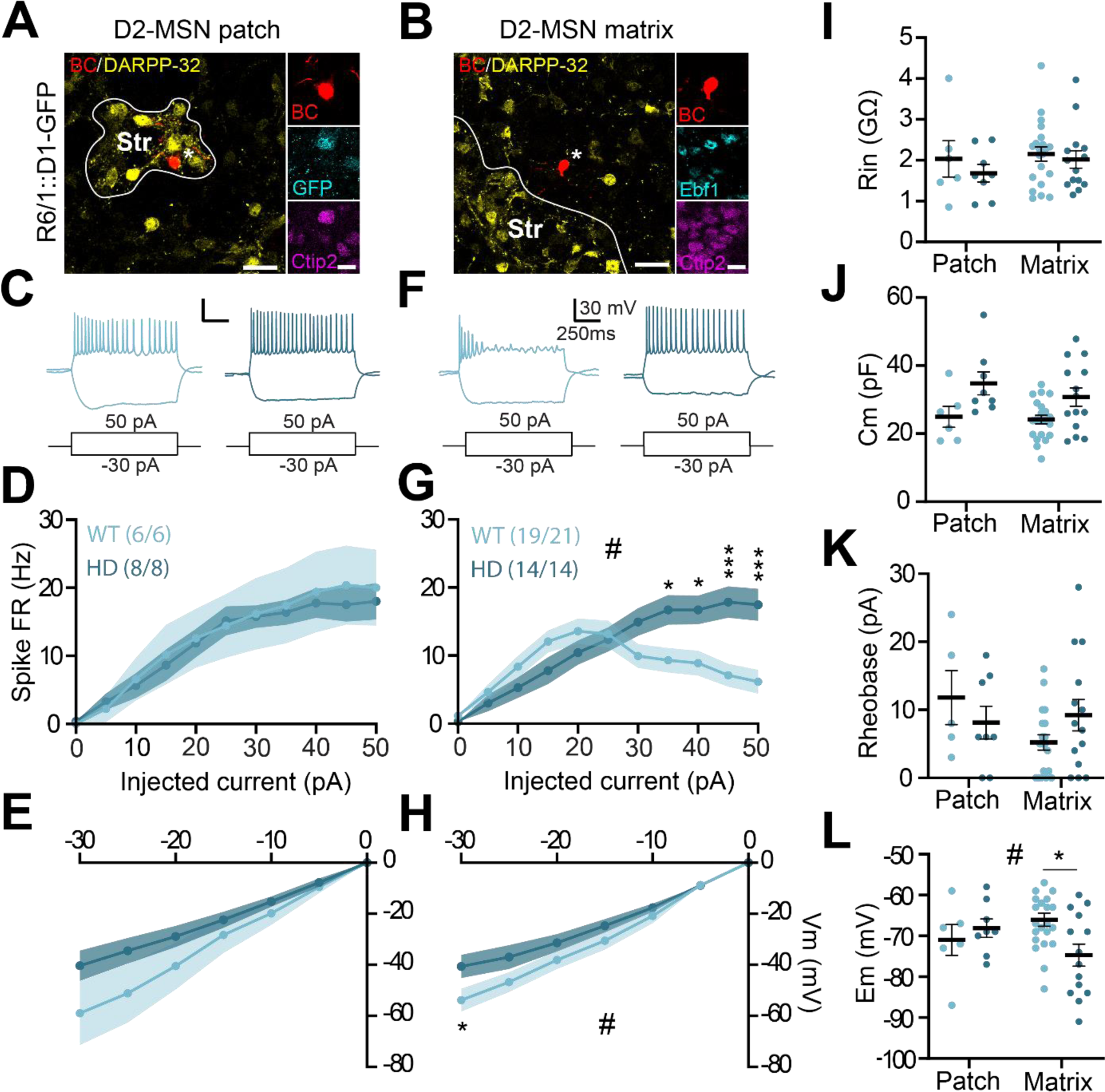
Electrophysiological properties of D2-MSNs from R6/1 mice are altered in a compartment-dependent manner during the first postnatal days. **A-B**. Examples of D2-MSNs neurons recorded in either striosome (**A**; GFP-/Ebf1+/DARPP32+) or matrix (**B**; GFP-/Ebf1-/DARPP32-). **C, F**. Representative D2-MSNs responses from striosome (**C**) and matrix (**F**) in WT (light blue) and R6/1 (dark blue) mice following the injection of hyperpolarizing and depolarizing currents. **D, G**. Mean discharge frequency/injected current curves obtained for each condition. **E, H**. Membrane potential/current curves obtained for each condition. **I-L**. Main electrophysiological properties of D2-MSNs showing input resistance (**I**), capacitance (**J**), rheobase (**K**) and resting membrane potential (**L**) as a function of the striatal compartment. Scale bars: 25 and 10 μm.

Upon distinguishing neurons by compartmental location, we found that alterations previously observed were not present in striosomal neurons. Specifically, no significant differences were observed in evoked discharge frequency or the injected potential/current curve in D2-MSNs from striosomes of R6/1 mice compared to WT (**Figure 3D, E**). Additionally, none of the recorded membrane parameters showed significant differences between striosomal D2-MSNs from R6/1 and WT mice (**Figure 3I-L**).

In contrast, matrix D2-MSNs from R6/1 mice exhibited exacerbated alterations compared to WT mice, consistent with previous observations. Specifically, the frequency of evoked discharge was significantly different for injected currents between 35 and 50 pA, as neurons from R6/1 mice were able to generate action potentials at higher injected currents compared to WT (**Figure 3F-G**). Hyperpolarization induced by current injection was significantly reduced in D2-MSNs from R6/1 mice at -30 pA, indicating accelerated expression of rectifier currents (**Figure 3H**). Additionally, the resting membrane potential of matrix D2-MSNs from R6/1 mice was significantly more hyperpolarized than that of WT mice (**Figure 3L**). We performed the same analysis for D1-MSNs and found no differences in the excitability of this cell-type between WT and R6/1 mice in both striosomal and matrix compartments (**Figure S5**). In summary, the observed maturation defects in the electrophysiological properties of D2-MSNs from R6/1 mice at P0-2 are specific to matrix neurons, with disturbances disappearing within the first postnatal week.

### The dendritic complexity of D2-MSNs appears to be altered during the early postnatal period in R6/1 mice

We examined the evolution of dendritic arborization of D2-MSNs by measuring its length and analyzing its complexity using Scholl analysis. Additionally, we measured other parameters such as the number of nodes (representing branches in the dendritic tree) and the number of primary dendrites. We observed significant maturation of dendritic arborization in D2-MSNs, with the dendritic length doubling from around 1 mm to 2.5 mm during the first two postnatal weeks in WT mice (**Figure 4A, C**). This indicates progressive extension of dendrites into a wider area of the striatum. Furthermore, the number of nodes increased progressively with age, reflecting an overall increase in dendritic complexity, while the number of primary dendrites remained constant.

**Figure 4:**
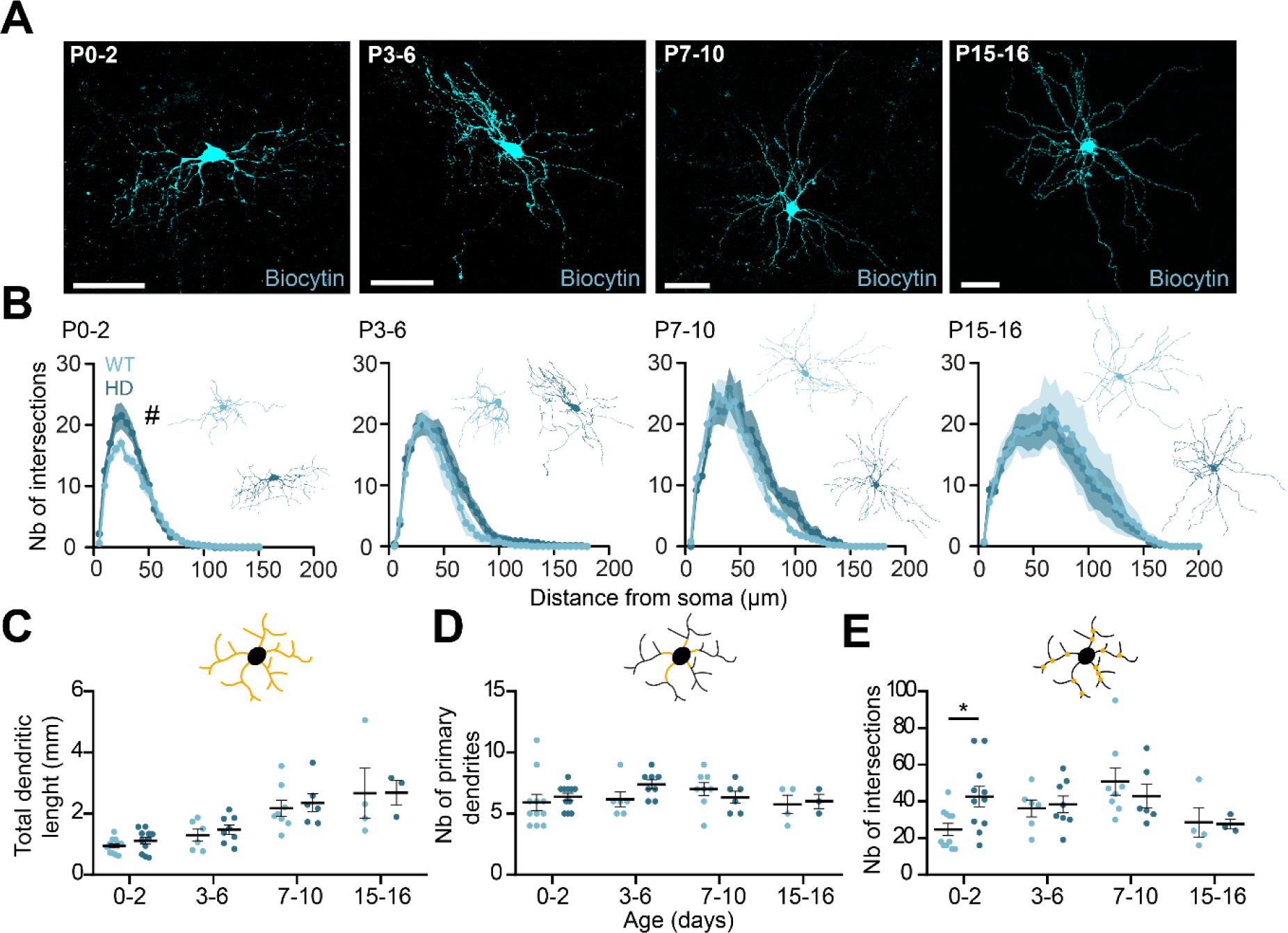
Maturation of the morphological properties of D2-MSNs is impaired in R6/1mice during the first postnatal days. **A.** Examples of biocytin-filled D2-MSNs at different postnatal stages. **B.** Scholl analysis illustrating the number of intersections between the dendritic tree and concentric circles spaced 5 μm apart from the soma. **C-E.** Main features of the dendritic arborization with dendritic length **(C)**, number of primary dendrites **(D)** and number of nodes **(E)**. Scale bars: 50 μm.

When comparing the evolution of dendritic length between WT and R6/1 mice, no significant differences were observed at any age (p>0.05; Student’s t test). However, changes in dendritic complexity were noted in D2-MSNs from R6/1 mice at P0-2. Scholl analysis revealed significant differences compared to WT mice (# F_(29, 609)_ = 2.856; p<0.0001; repeated measures ANOVA), with an overall increase in the number of intersections in R6/1 mice (**Figure 4B**). However, the shape of the curve was not shifted to the right. The total dendritic length (**Figure 4C**) and the number of primary dendrites (**Figure 4D**) were not different between the two genotypes. However, the number of dendritic nodes appeared to be significantly increased in R6/1 mice at P0-2 compared with WT mice (**Figure 4E**). We performed the same morphological analysis on D1-MSNs and, conversely to D2-MSNs, no difference was observed between WT and R6/1 mice (**Figure S6**). Together, these observations suggest that D2-MSNs from R6/1 mice develop abnormal dendritic branching without extending further into the striatum during the early postnatal period.

### Cortico-striatal synaptic transmission is altered in D2-MSNs during the early postnatal period in R6/1 mice

It is known in the literature that synaptic excitatory afferents are rapidly established after birth and continue to develop during the first postnatal weeks (Dehorter et al., 2011; Krajeski et al., 2019). To this end, we first measured spontaneous excitatory postsynaptic currents (sEPSCs) using whole-cell *ex vivo* electrophysiological recordings at three different developmental stages (P0-2, P3-6 and P7-10; **Figure 5A-B**). sEPSCs were recorded at a holding potential of -70 mV and in presence of GABA_A_ and GABA_B_ receptors antagonists (5µM GBZ + 1µM CGP55845) in the ACSF solution perfused into the recording chamber (**Figure 5A-B**). The glutamatergic nature of the recorded currents was confirmed by the addition of AMPA and NMDA receptor antagonists (20µM DNQX + 50µM D-APV), which resulted in the complete disappearance of spontaneous EPSCs (**Figure 5C**). The frequency of the currents increased progressively with age (WT: P0-2: Fq = 0.49 ± 0.13 Hz, P3-6: Fq = 0.63 ± 0.12 Hz, P7-10: Fq = 0.84 ± 0. 15 Hz; **Figure 5D**), while the mean amplitude of EPSCs remained stable (WT: P0-2: Amplitude = 17.69 ± 1.79 pA, P3-6: Amplitude = 14.99 ± 1.75 pA, P7-10: Amplitude = 16.22 ± 1.87 pA; **Figure 5E**). Comparing the EPSC frequency values between the two genotypes, we observed no significant difference, at any age (p>0.05, Student’s t test; **Figure 5D**). Similarly, between P0 and P6, there was no significant difference in the amplitude of the EPSCs (P>0.05, Student’s t test). However, from P7 onwards, amplitude values were significantly reduced in R6/1 mice (WT: mean = 16.22 ± 1.87 pA; R6/1: mean = 11.01 ± 1.01 pA; P=0.0299, Student’s t

**Figure 5:**
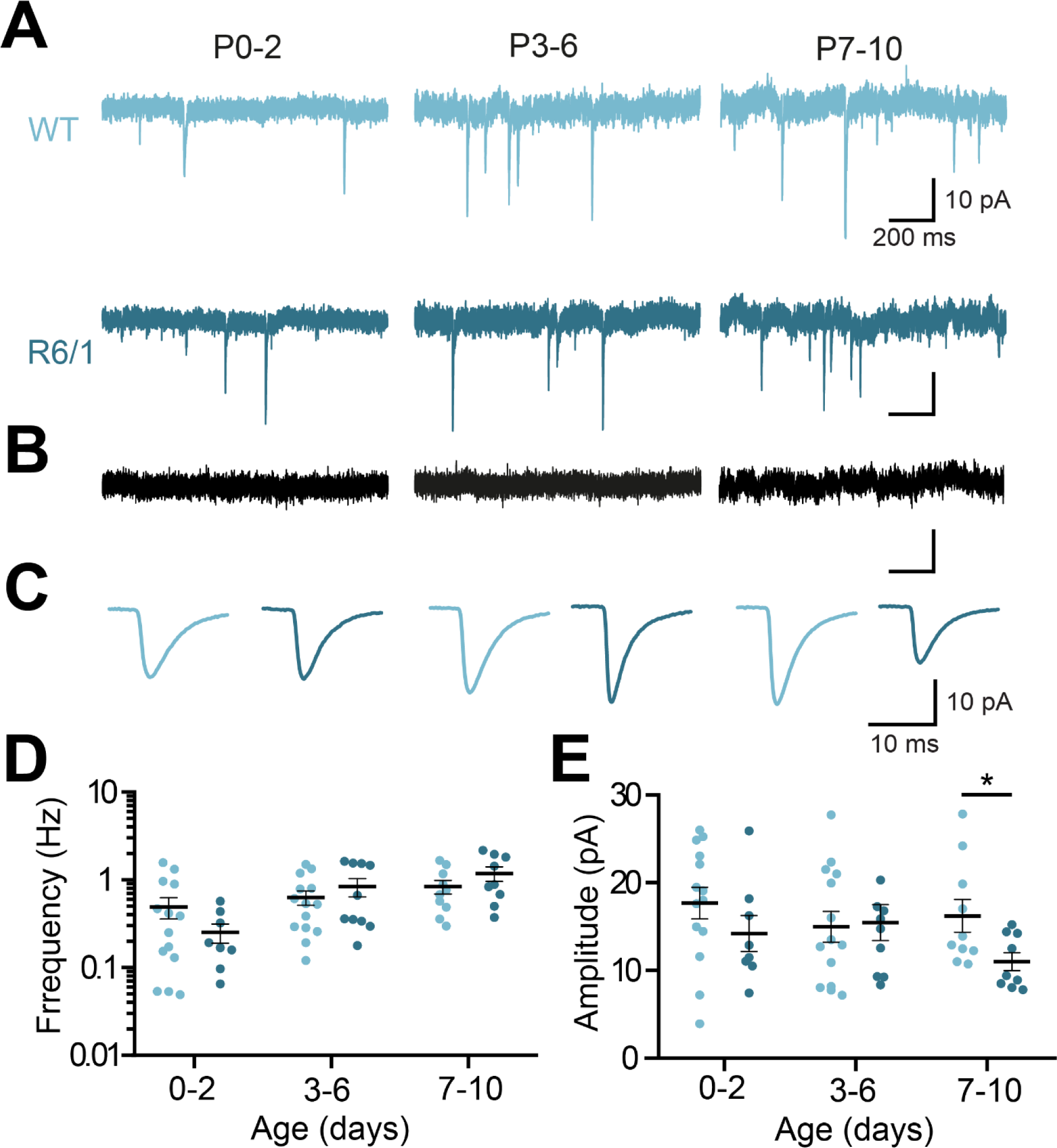
Glutamatergic transmission is impaired in D2-MSNs from R6/1 mice during the second postnatal week. **A**. Representative recordings of spontaneous excitatory postsynaptic currents (sEPSCs) imposed at a potential of -70 mV in WT (top, light blue) and R6/1 (bottom, dark blue) mice at P0-2, P3-6 and P7-10, recorded in presence of GABA_A_ and GABA_B_ antagonists (5 µM GABAzine + 1µM CGP55485). **B**. sEPSCs disappear when AMPA and NMDA antagonists (20µM DNQX + 50µM APV) are added to the bath. **C**. Average traces of sEPSCs obtained for each condition. **D-E**. Mean frequency (**D**) and amplitude (**E**) of sEPSCs as a function of age.

test; **Figure 5E**). sEPSCs were also recorded in D1-MSNs (**Figure S7**) under the same experimental conditions. When comparing sEPSCs frequencies we observed a significant reduction (WT: P7-10: Fq = 0.76 ± 0.14 Hz; R6/1: P7-10: Fq = 0.37 ± 0.06 Hz, P=0.0155, Student’s t-test; **Figure S7D**) while the amplitude of sEPSCs remained similar across the three developmental ages between the two genotypes (p>0.05, Student’s t-test; **Figure S7E**). We also measured sEPSCs in D2-MSNs in CAG140 mice (**Figure S8**), but did not observe any significant change between the genotypes. Given that changes in excitability observed in CAG140 mice occured later than in R6/1 mice, it would be pertinent to study synaptic transmission after P10.

Previous results have demonstrated alterations in glutamatergic transmission in the striatum during the second postnatal week. The cortex and thalamus represent the two main sources of glutamatergic afferents to the striatum (Y. Smith et al., 2004; Voorn et al., 2004). Given the particular vulnerability of the cortico-striatal pathway in HD (Blumenstock & Dudanova, 2020), we decided to study specifically this pathway. To this end, we performed whole-cell patch-clamp recordings of D2-MSNs in the dorsal striatum and electrically stimulated cortical afferents via a bipolar electrode placed at the level of the corpus callosum (**Figure 6**). These experiments were carried out in the presence of GABAergic antagonists to avoid the recruitment of inhibitory afferents. We specifically performed these recordings at P7-10, as it was at this age that we observed differences in the amplitude of the sEPSCs measured (**Figure 5**). First, we performed a double stimulation protocol at 20 Hz to measure the paired-pulse ratio (PPR). When we compared the PPR between WT and R6/1 mice, we found no significant difference, suggesting no change in the short-term release probability of glutamate at cortico-striatal synapses on D2-MSNs in R6/1 mice (**Figure 6A-B**). Next, we examined the contribution of AMPA and NMDA receptors to cortico-striatal transmission. To do this, we recorded evoked synaptic currents either at -70 mV to specifically observe AMPA receptor-mediated currents, or at +80 mV to record currents mediated by both AMPA and NMDA receptors. As AMPA receptors exhibit fast kinetics and inactivate much faster than NMDA receptors, we measured the amplitude of currents 50 ms after stimulation to consider only the NMDA component of the EPSC (**Figure 6C**). Measuring the ratio between the amplitude values of NMDA and AMPA responses, we found that this ratio was significantly increased in R6/1 mice compared with WT mice (WT: NMDA/AMPA = 0.81 ± 0.17, R6/1: NMDA/AMPA = 1.71 ± 0.31; **p=0.0014,

**Figure 6:**
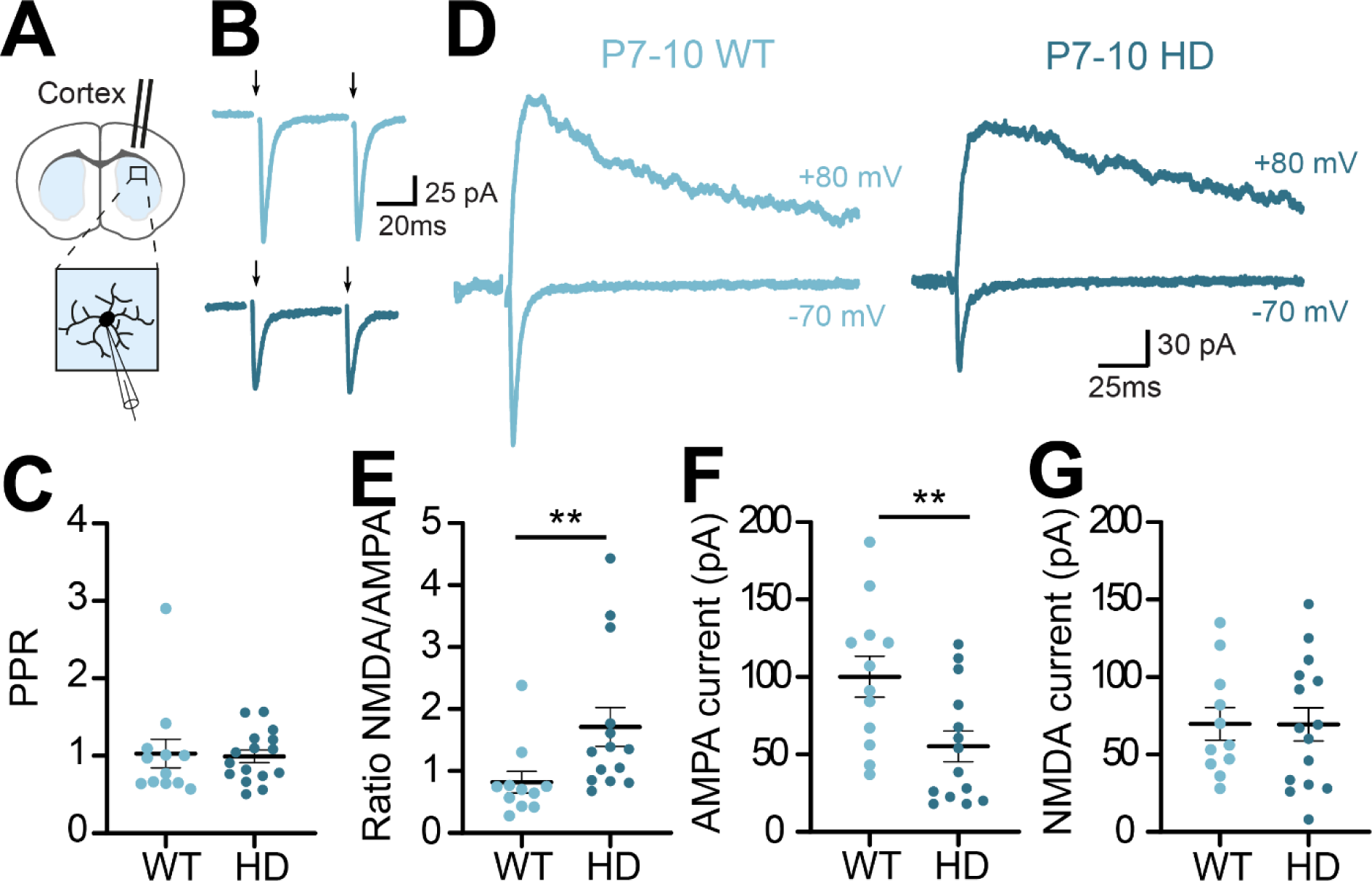
Cortico-striatal transmission in D2-MSNs is impaired in R6/1 mice during the second postnatal week. **A.** Schematic of the experiment showing the position of the stimulating electrode and the recording electrode. **B.** Representative recordings of evoked excitatory postsynaptic currents (eEPSCs) at a potential of -70 mV in WT (light blue) and R6/1 (dark blue) mice. **C.** Average paired pulse ratio (EPSC2/EPSC1). **D.** Representative recordings at an imposed potential of either -70 mV to record AMPA receptor-mediated currents or at +80 mV to record currents mediated by both AMPA and NMDA receptors. **E.** NMDA/AMPA ratio obtained by dividing the amplitude values of NMDA currents by those of AMPA currents. **F-G.** Amplitude values of AMPA (F) and NMDA (G) currents. All recordings were performed in presence of GABA_A_ and GABA_B_ receptors antagonists (5 µM GABAzine + 1 µM CGP55845).

Mann-Whitney U test; **Figure 6D**). Upon examining the currents mediated by AMPA receptors (**Figure 6E**) and NMDA receptors (**Figure 6F**), we found that this change in ratio was caused by a significant decrease in AMPA currents (WT: AMPA amplitude = 100. 2 ± 13.3 pA, R6/1: AMPA amplitude = 55.2 ± 9.9 pA, **p=0.0088, Mann-Whitney test) with no change in the amplitude of NMDA responses (WT: NMDA amplitude = 69.7 ± 10.5 pA, R6/1: NMDA amplitude = 69.4 ± 10.7 pA, P=0.98, Student’s t test). These observations thus suggest a weakening of cortico-striatal transmission in D2-MSNs in R6/1 mice during the second postnatal week.

We also studied cortico-striatal transmission in D1-MSNs (**Figure S9**). When we measured PPR in D1-MSNs from WT and R6/1 mice, we found no significant difference between the two genotypes (p>0.05, Mann-Whitney test; **Figure S9B**). Furthermore, when measuring the NMDA/AMPA ratio, we observed no significant differences between genotypes (WT: NMDA/AMPA = 1.41 ± 0.24, R6/1: NMDA/AMPA = 1.92 ± 0.50, p>0.05, Student’s t test; **Figure S9C-D**), nor were there any changes in the specific amplitudes of AMPA or NMDA currents (**Figure S9E-F**).

In conclusion, these experiments revealed a specific alteration in glutamatergic transmission in D2-MSNs during the second postnatal week in R6/1 mice, which seems to be due to a reduction in postsynaptic AMPA receptor currents.

## Discussion

Our work demonstrates that the electrical properties and morphology of D2-MSNs are transiently modified from birth in the striatum of HD mice. In addition, we have shown a reduction in cortico-striatal glutamatergic transmission in D2-MSNs starting from the second postnatal week. It is well-established that the development of the cortex and striatum is closely linked during the postnatal stage, since alterations in either cortical or striatal activity during this period induces significant changes in cortico-striatal connectivity (Kozorovitskiy et al., 2012; Peixoto et al., 2016). Moreover, it appears that modifications of early neuronal activity, whether an increase or a decrease, have an impact on the formation of a functional adult circuit (Bitzenhofer et al., 2021; Braz et al., 2022). This notion has been notably highlighted in several models of neurodevelopmental diseases such as fragile X syndrome, schizophrenia, and autism spectrum disorders (Ahmed et al., 2023; He et al., 2019; Kirischuk et al., 2017).

### General observations

During the first postnatal week, the percentage of GFP^+^ neurons in the striatum of D1-GFP mice was very low, mainly located in MOR^+^ striosomes and gradually colonizing the matrix over the second postnatal week. This observation had already been made in D1-GFP mice at P0, but no precise quantification of the evolution of D1 receptor expression had been carried out (Merchan-Sala et al., 2017). From only 4% of MSNs expressing GFP at P0, this percentage gradually increases to reach 42% at P15. This agrees with various studies showing that the expression of the Drd1 gene is low at embryonic and early postnatal stages compared to adult stages (Araki et al., 2007; Caille et al., 1995; Jung & Bennett, 1996; Rao et al., 1991; Yang et al., 2021). We performed the same analysis in D2-GFP mice, where the proportion of GFP^+^ D2-MSNs is close to 50% from P0 (40%) and remains stable over time, consistent with the proportions of D1-MSNs/D2-MSNs observed in adults (Gagnon et al., 2017). These results indicate that during the first postnatal days, there is a significant imbalance between the expression of D1 and D2 receptors. Interestingly, a study showed that D1 and D2 receptors play opposite roles in neuronal migration, with D1 receptors promoting migration while D2 receptors seem to inhibit it (Crandall et al., 2007). The imbalance in favor of D2 receptor expression during the first postnatal days could contribute to the stabilization of the circuit once the neurons have migrated and reached their final destination. It is of note that the distribution of D1-and D2-MSNs is not altered in the striatum of the R6/1 mice during the first postnatal 15 days.

### mHtt impacts the electrophysiological and morphological properties of matrix D2-MSNs

The postnatal period is marked by the progressive maturation of the electrophysiological properties of MSNs, as they acquire characteristics like those of adult neurons (Dehorter et al., 2011; Krajeski et al., 2019; Tepper & Trent, 1993). Comparing this maturation between WT and R6/1 mice, we observed no alterations in D1-MSNs. In contrast, D2-MSNs showed abnormal hyperpolarization and increased capacitance, reflecting a decrease in excitability during the first postnatal days. Moreover, an inward rectifying current seems to be already present in these neurons, suggesting an earlier acquisition of the Kir channels involved in establishing the highly hyperpolarized resting potential of MSNs (Kita et al., 1984; Nisenbaum et al., 1996) and could therefore explain this early hyperpolarization of D2-MSNs. These alterations are transient and limited to the first postnatal days, suggesting the development of compensatory mechanisms aimed at restoring normal circuit function. Interestingly, transient alterations in electrophysiological properties and dendritic arborization of pyramidal neurons have also been observed during the first weeks after birth in the cortex of HD mice (Braz et al., 2022).

The alterations observed in R6/1 mice were specific to D2-MSNs within the striatal matrix. It is known that striosomal and matrix neurons have different afferents and efferents and are thus involved in distinct functions (Crittenden & Graybiel, 2011; Flaherty & Graybiel, 1994; Friedman et al., 2015). Moreover, matrix MSNs predominantly establish their axonal connections with their target structures during the first postnatal week (Fishell & van der Kooy, 1989, 1991; Krajeski et al., 2019). In HD, these different compartments are associated with the different symptoms observed; striosomes contribute to the appearance of psychiatric symptoms while the matrix is involved in the appearance of motor symptoms (Matsushima et al., 2023; Morton et al., 1993). Thus, it is noteworthy that specific developmental alterations occur during a critical period for circuitry integration and within the subpopulation of MSNs that will subsequently be involved in the appearance of major motor symptoms in adulthood.

### mHtt impacts glutamatergic transmission on D2-MSNs

Although cortical and thalamic afferents are partially present from birth, cortico-striatal synaptogenesis continues throughout the postnatal period (Dehorter et al., 2011; Krajeski et al., 2019; Peixoto et al., 2016). We observed a decrease in the amplitude of spontaneous excitatory postsynaptic currents (sEPSCs) in D2-MSNs of R6/1 mice from P7. As the cortico-striatal pathway is particularly altered in adulthood (Blumenstock & Dudanova, 2020), we specifically studied the responses evoked by electrical stimulation of cortico-striatal axons at P7-10. We observed a significant increase in the NMDA/AMPA ratio in D2-MSNs. This change in ratio is caused by a decrease in the amplitude of the currents induced by AMPA receptors, without a change in the currents induced by NMDA receptors suggesting a postsynaptic reduction in AMPA receptors. Interestingly, similar observation has been made in the developing cortex of HQ111 mice (Braz et al., 2022). Taken together, our results indicate that mHtt induces a postsynaptic decrease in cortico-striatal glutamatergic transmission in D2-MSNs of HD mice during the second postnatal week. Several mechanisms can contribute to the reduction in AMPA currents in HD mice, including alterations in AMPA receptor transport and mobility on the surface, a reduction in receptor expression or phosphorylation. It is already known that in adulthood, mHtt reduces glutamatergic transmission by impacting the trafficking and surface diffusion of these receptors (Mandal et al., 2011; Zhang et al., 2018). Interestingly, treating HD mice with AMPAkine, a positive allosteric modulator of AMPA receptors, has been shown to not only restore the observed alterations but also to delay the appearance of major HD symptoms in adulthood (Braz et al., 2022). This suggests a critical role of glutamatergic synaptic transmission in the early dysfunction of the corticostriatal microcircuit in HD.

It is remarkable that alterations are preferentially observed in D2-MSNs, both during development and in adulthood, even though mHtt is ubiquitously expressed in both MSN subpopulations (Albin et al., 1992; Deng et al., 2004; Reiner et al., 1988). In adulthood, decreased BDNF release at cortico-striatal synapses and decreased TrkB receptor expression in striatal neurons are known to play a crucial role in HD-related neurodegeneration (Ginés et al., 2006; Saudou & Humbert, 2016). Interestingly, it appears that the TrkB receptor directly influences potassium channel activation (Carrillo-Reid et al., 2019; Rogalski et al., 2000), and this phenomenon has been shown to be impaired in a mouse model of HD (Carrillo-Reid et al., 2019). Moreover, it has been observed that TrkB receptor expression is predominantly found in D2-MSNs, with 98% of D2-MSNs expressing TrkB at P10, compared with only 18% of D1-MSNs (Baydyuk et al., 2011; Baydyuk & Xu, 2014). Although it is not known whether TrkB receptor expression is reduced during development in the presence of mHtt, the selective expression of this receptor in D2-MSNs could explain the specificity of the alterations we observed. All these observations lead us to hypothesize that mHtt could result in a decrease in TrkB receptor expression in D2-MSNs, leading to an early expression of potassium conductances, thus specifically disrupting D2-MSN activity during the postnatal period.

In conclusion, our data demonstrating postnatal alterations specifically in D2-MSNs are consistent with existing literature suggesting a preferential vulnerability of these neurons in HD (Albin et al., 1992; Deng et al., 2004; Reiner et al., 1988). Additionally, our findings align with the notion that the development of the cortex and striatum is closely linked during the postnatal stage, as alterations in either cortical or striatal activity can induce significant changes in cortico-striatal connectivity (Kozorovitskiy et al., 2012; Peixoto et al., 2016). Our results suggest that the initial alterations in excitability and morphology in D2-MSNs may influence the cortico-striatal transmission received by these neurons. Therefore, it appears that early alterations in the striatal circuit, particularly in D2-MSNs, could play a pivotal role in the vulnerability of these neurons and in the progression of HD. These findings contribute to our understanding of the pathophysiology of HD and may offer insights for developing targeted therapeutic interventions aimed at preserving D2-MSNs function and mitigating disease progression.

## Author contributions

M.L., M.G. and J. B. conceived and designed the experiments. M.L. M.G. L.B., performed and analyzed the anatomical and neuronal reconstruction studies. G.C. develop the scripts to quantify neuronal density. M.L, J.S. and Q. R. performed and analyzed the ex vivo electrophysiology experiments. M.L., M. G. and J.B. wrote the manuscript and M.L. prepared the figures.

## Acknowledgments

M.L. received a Ph. D fellowship from the Association Huntington France. This work was supported by recurrent funding from the University of Bordeaux and the CNRS. This study also received financial support from the French government in the framework of the University of Bordeaux’s IdEx “Investments for the Future” program / GPR BRAIN_2030.

## Experimental model and subject details

### Animals

Experiments were performed in two mouse models of HD: a model expressing a truncated form of the human huntingtin gene, the R6/1 model, and a knock-in CAG140 model. R6/1::D1-GFP mice were generated by crossing heterozygous R6/1 male with homozygous D1-GFP females. Resultant WT::D1-GFP or R6/1::D1-GFP mice were obtained in half proportion. R6/1::D2-GFP or CAG140::D2-GFP mice were generated by crossing heterozygous R6/1- or CAG140 males with homozygous D2-GFP females These mice lines reveal D1- or D2-MSNs by the presence of green fluorescent protein. All animals were maintained in a 12/12h light/dark cycle, in stable conditions of temperature and humidity, with access to food and water *ad libitum*. We verified the CAG repeat number in our R6/1 and CAG 140 lineages. Our R6/1 lineage exhibited a reduction in the CAG repeat number compared to the norm, with an average of 55 repeats instead of the expected 115 (Mangiarini et al., 1996). This extremely rare contraction has been described in the literature and leads from 1 year of age to mHtt aggregates similar to that found in human HD (Morton et al., 2019). The extension repeat number in our CAG140 lineage was 109. All experiments were approved by the local ethical committees and by the French Ministry of Research and APAFIS#14255.

### Tissue preparation and immunohistochemistry

Mice (0-15 days) were deeply anaesthetized with isoflurane and then decapitated. Brains were extracted and post-fixed overnight in a solution of 4% w/v PFA in 0.1 M phosphate buffered saline, pH 7.4 (PBS) at 4C. Next day, brains were washed twice in PBS and then stored in PBS-azide 0,03% before being cut into 50 μm coronal sections on a vibratome (VT1000S; Leica Microsystems). For immunohistochemistry, sections were washed 3 times in PBS for 10 minutes. Sections were then incubated in blocking solution containing 4% Normal Donkey Serum (NDS) in PBS with 0,3% Triton X-100 for 1 hour at room temperature with constant shaking. Finally, sections were incubated in primary antibodies diluted in 4% NDS in PBS with 0,3% Triton X-100 solution overnight at 4°C. The following antibodies were used: rat anti-Ctip2 (1:1000, Abcam ab18465) and chicken anti-GFP (1:1000, Aves Lab GFP1010). Ctip2 is expressed by all MSNs but not interneurons (Arlotta et al., 2008). Next day, sections were rinsed three times in PBS for 10 min before incubation for two hours at room temperature with secondary antibodies. The following fluorescent probes were used: donkey anti-rat Alexa Fluor 647 (1:500, Jackson Laboratories), donkey anti-chicken Alexa Fluor 488 (1:500, Jackson Laboratories). Sections were rinsed 3 times for 10 min in PBS before mounting in ProLong Diamond Antifade Mountant (Invitrogen).

### Image acquisition, analysis and quantification

Fluorescence images were acquired either on a confocal microscope (Carl Zeiss© LSM 900) using ZEN software or on a spinning disk microscope using Metamorph software (Leica DMI6000B). For cell density analysis, three rostro-caudal levels of the striatum were defined using the anterior commissure as an anatomical landmark. A tilescan of the whole striatum on a single plane was then acquired using a x40 or x20 objectives. For cell density analysis, we manually delineated each striatal area to calculate the specific area of each level. Neuronal counting was then performed semi-automatically using the QuPath software (Bankhead et al., 2017). To determine the density of the different neuronal populations, the number of neurons identified in each striatum was divided by the area of that striatum. The density of D1-MSNs or D2-MSNs was measured by dividing the number of GFP^-^ or GFP^+^ neurons, respectively, by the area of the striatum.

### Acute brain slices preparation and recording conditions

Coronal striatal slices were prepared from 0-16-day-old R6/1, CAG140 or non-carrier control mice. Briefly, animals were deeply anaesthetized with isoflurane and decapitated. The brains were removed from the skull and placed in ice-cold modified artificial CSF (aCSF), saturated with 95% O_2_ and 5% CO_2_, and containing (in mM): 230 sucrose, 10 MgSO4·7H2O, 2.5 KCl, 1.25 NaH2PO4·H2O, 0.5 CaCl2·H2O, 26 NaHCO3, et 10 D-glucose. Brains were sectioned into 300 µm-thick coronal slices by using a vibratome (VT-1200S; Leica Microsystems, Germany). Slices were then immediately transferred in ACSF containing (in mM): 126 NaCl, 2.5 KCl, 1.25 NaH2PO4·H2O, 2 CaCl2·H2O, 2 MgSO4·7H2O, 26 NaHCO3, 10 D-glucose, 5 L-glutathion and 1 sodium pyruvate (gassed with 95% O_2_/5% CO_2_). Slices were incubated 1 hour at 32°C in this solution and then maintained at room temperature in the same solution until recording. Striatal slices were transferred to a recording chamber, continuously perfused with oxygenated ACSF containing (in mM): 126 NaCl, 3 KCl, 1.25 NaH2PO4·H2O, 1.6 CaCl2·H2O, 1.5 MgSO4·7H2O, 26 NaHCO3, and 10 D-glucose (at 32°C with a perfusion speed of 2 mL.min^-1^). Striatal neurons were visualized using infrared gradient contrast video microscopy (E600FN, Eclipse workstation, Nikon, Japan) and a water-immersion objective (Nikon Fluor 60 X/1.0 NA). MSNs were distinguished from interneurons by their morphology and GFP-expressing neurons were identified in real time with an epifluorescence (Nikon Intensilight C-HGFI). Recordings from individual neurons were made using patch pipettes (5–8 MΩ) pulled from thick-walled borosilicate glass capillaries (G150–4; Warner Instruments, Hamden, CT, USA) on a micropipette puller (P-97, Sutter Instruments, Novato, CA, USA). For recordings of the intrinsic properties, the patch pipette internal solution contained (in mM): 135 K-gluconate, 3.8 NaCl, 1 MgCl2·6H_2_O, 10 HEPES, 0.1 Na_4_EGTA, 0.4 Na2GTP, 2 MgATP and 5.3 biocytin. The osmolarity and pH of the intrapipette solution were adjusted at 290 mOsm and 7.2 respectively. In current-clamp mode, the junction potential (+13mV; JPCalc, Clampex 10) was corrected. For recordings of cortico-striatal transmission, the patch pipette internal solution contained (in mM): 132 CsMSO_3_, 3.6 NaGlu, 1 MgCl2·6H_2_O, 10 HEPES, 0.1 Na_4_EGTA, 5 TEA-Cl, 0.4 Na2GTP, 2 MgATP, 5 QX-314 and 5.3 biocytin. The recordings were amplified using a Multiclamp 700B amplifier (Molecular Devices) and digitized at 20 kHz (Digidata 1320A/1550B) using Clampex 10 acquisition software (Molecular Devices, Sunnyvale, CA, USA).

### Stimulation and recording protocols

In voltage-clamp mode, the holding potential was set at -60 mV and successive voltage steps of -5 mV were performed to measure passive properties of the cells including input resistance and capacitance. In current-clamp mode, hyperpolarizing and depolarizing current steps were injected through the patch pipette to assess the active properties of MSNs (rheobase and spiking frequency). Current steps ranges were for P0-2: -30 to +50 pA; for P3-6: -50 to +50 pA; for P7-10: -50 to +80 pA and P15-16: - 200 to +250 pA, 1000-ms duration. Spontaneous post-synaptic currents (EPSCs) were recorded in voltage-clamp mode at a fixed membrane potential of -70 mV during 5 and 10 minutes. For these experiments, recordings were conducted in the presence of the GABA_A_ antagonist SR95531/GABAzine (5 µM, Tocris Bioscience) and the GABA_B_ antagonist CGP55845 (1 µM, Tocris Bioscience). To confirm the glutamatergic nature of the measured currents, antagonists of AMPA receptors (DNQX, 20 µM, Abcam) and NMDA receptors (D-AP5, 50 µM, Abcam) were used in some experiments. All pharmacological agents used were prepared as stock solutions, aliquoted in distilled water, and stored at -20°C. They were then diluted to experimental concentrations on the day of the experiment and perfused into the recording chamber. Activation of cortical afferents was achieved using a bipolar stimulation electrode placed just above the corpus callosum (stimulation intensity between 100 and 800 µA). These experiments were conducted in the presence of GABAergic transmission blockers (5 µM GABAZine and 1 µM CGP55845). Double stimulations of 0.1 ms at 5 and 20 Hz were performed to measure the amplitude of EPSCs and assess the short-term plasticity of the cortico-striatal synapse determined through the paired-pulse ratio (PPR=Pulse2/Pulse1). For these experiments, neurons were recorded at a potential of -70 mV. To determine the relative contribution of AMPA/Kainate and NMDA receptors in cortico-striatal transmission in MSNs, synaptic responses to single axonal stimulation of cortico-striatal neurons were recorded at -70 mV and +80 mV, respectively.

### Analysis of recordings

Data analyses were performed using the pClamp 10.6 and OriginPro 7.0 software (Microcal Software). Series resistance (Rs), membrane resistance (Rm) and membrane capacitance (Cm) were calculated in whole-cell voltage-clamp configuration by analysis of the transient currents in response to a -5 mV step of potential (ΔV). Cells with Rs exceeding 30 MΩ were excluded from analysis. The Rm was measured using Ohm’s law (U = RI) based on the steady-state current at the end of the potential step. The Cm was calculated using the following formula: Cm = Q/ΔV in which Q correspond to the area under the capacitive transient after removing leak currents. In current-clamp mode, action potential firing frequency and membrane potential were calculated for each injected current and F-I curve (frequency of action potential firing as a function of injected current) and I-V curve (membrane potential as a function of injected current) were constructed. The rheobase was also determined with progressive current steps of 10 ms. Electrically-evoked EPSCs amplitude were measured as the peak of inward current relative to the baseline holding current preceding the electrical pulse. To obtain the paired pulse ratio, the amplitude of the second EPSC was divided by the amplitude of the first EPSC (EPSC_2_/EPSC_1_). The AMPA/NMDA ratio was calculated from recordings of the same neuron held at -70 mV to observe AMPA currents or at +80 mV to observe NMDA currents. The AMPA receptor-mediated current corresponds to the amplitude of the peak from neuron held at -70 mV. The NMDA receptor-mediated current corresponds to the amplitude of the current, from neuron held at +80 mV, 50 ms after the electrical stimulation. Peak amplitude of AMPA receptor-mediated current was then divided by peak amplitude of NMDA receptor-mediated current to obtain the AMPA/NMDA ratio.

### Cell identification and cell reconstruction

Following whole-cell electrophysiology, the striatal slices were fixed in 4% paraformaldehyde in PBS. Subsequently, sections were processed essentially as described above using rat anti-Ctip2 and Chicken anti-GFP. In addition, the following antibodies were used: rabbit anti-Ebf1 (1:500, Millipore AB10523), rabbit anti PPE (1:2000, LsBio LS-C23084), Rabbit anti-MOR (1:1000, Immunostar 24216), Mouse anti-DARPP-32 (1:500, BD Biosciences 611520). Ebf1 and PPE antibodies staining were improved through antigen retrieval by heating sections at respectively 65°C for 1 hour and 80°C for 35 minutes in 10 mM sodium citrate (Target Retrieval Solution 10X, S1699; Aligent Technologies, Santa Clara (US); pH 6.1). Ebf1 and PPE are specific markers of respectively D1-MSNs (Garel et al., 1997) and D2-MSNs (Krajeski et al., 2019; Lee et al., 1997; Sharott et al., 2017). As Ebf1 is a transcription factor that progressively disappears during the first postnatal week, Ebf1 labelling was used to identify D1-MSNs from P0 to P4. PPE labelling was used to identify D2-MSNs from P5. Biocytin was revealed with Streptavidin-AF568 (1:2000, Invitrogen S11226).

Fluorescence images were acquired using a confocal microscope (Carl Zeiss© LSM 900) utilizing the ZEN software. For neuron identification, z-stack acquisitions with a 2 μm step size were performed using a x40 oil immersion objective. The images were then overlap using ImageJ (NIH) to determine the nature of the recorded MSNs. Additionally, z-stack acquisitions with a 0.15-micron step size were obtained using a x40 objective to image the entire dendritic arborization of the neurons. Once imaged, neurons were then reconstructed and analyzed using Neurolucida and Neuroexplorer software (MBF Bioscience, Delft, The Netherlands). Only neurons presenting a complete dendritic tree were included in the analysis. Scholl analysis was generated by calculating the number of intersections between the dendritic tree of neurons and each concentric circle spaced every 5 µm from the soma. Dendritic length, number of nodes, and number of primary dendrites were also measured.

### Statistics

Statistical analysis was conducted using Prism 8 (GraphPad Software, USA). Data in the text are presented as mean ± standard error of the mean (SEM). For comparisons between two independent groups, an unpaired Student’s t-test was performed if the data followed a normal distribution. Otherwise, a non-parametric Mann-Whitney test was used. For comparison of groups or F-I response curves, statistical analysis was conducted using two-way analysis of variance (ANOVA), followed by Sidak’s multiple comparisons post hoc test. Data were considered significant for p < 0.05.

## Supplemental figures

**Figure S1:**
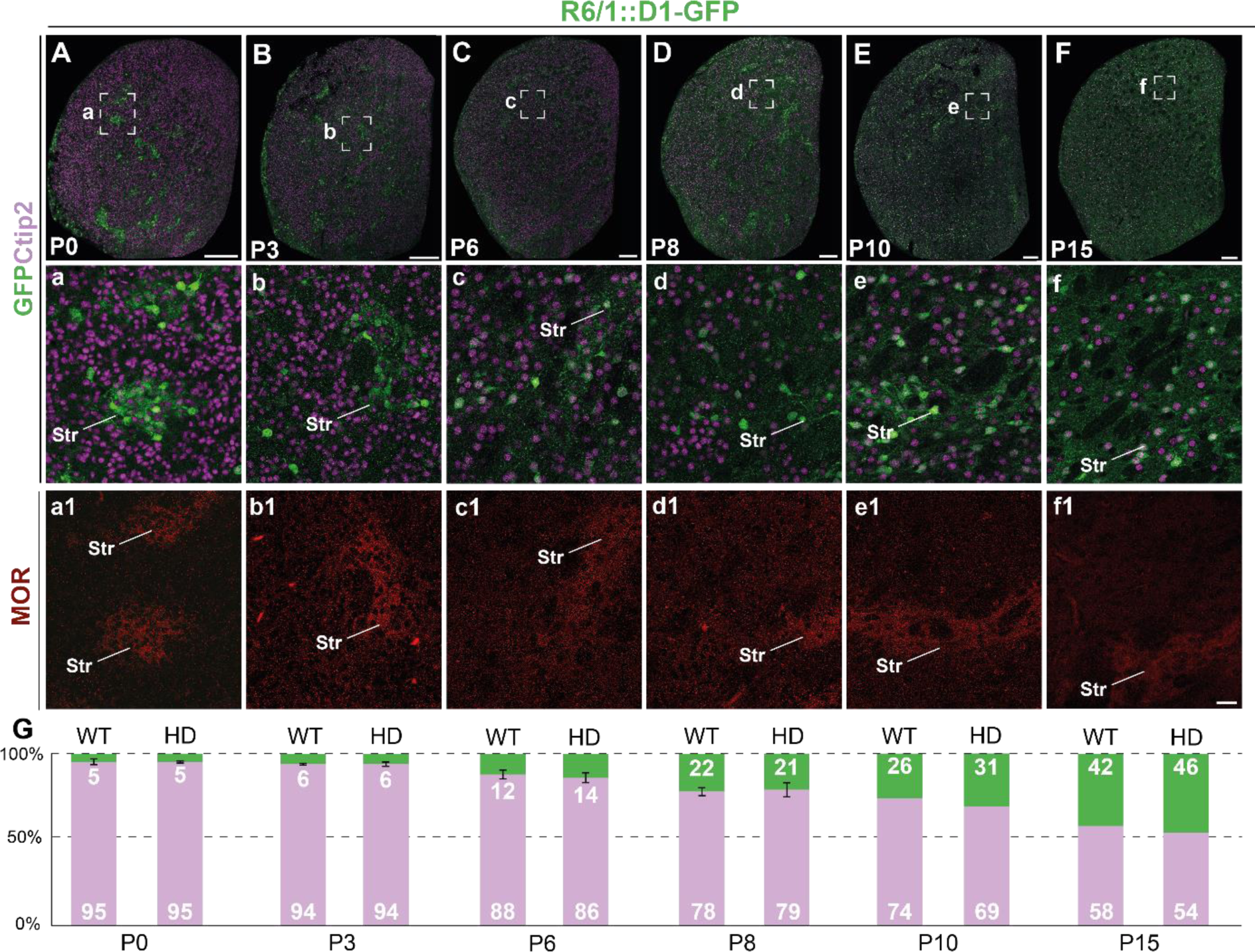
Distribution of GFP^+^ neurons in the striatum of WT::- and R6/1::D1-GFP mice during the first two postnatal weeks. **A-F.** Representative images of GFP and Ctip2 labeling at 6 developmental stages and enlarged images in a-f. a1-f1. µ-opioid receptor (MOR) labeling to distinguish striosomes. G. Quantification of the percentage of GFP^+^ D1-MSNs over the total number of Ctip2^+^ MSNs for WT and R6/1 mice at 6 developmental stages. Str: Striosomes. Scale bars: A-F, 200 μm; a-f1, 25 μm.

**Figure S2:**
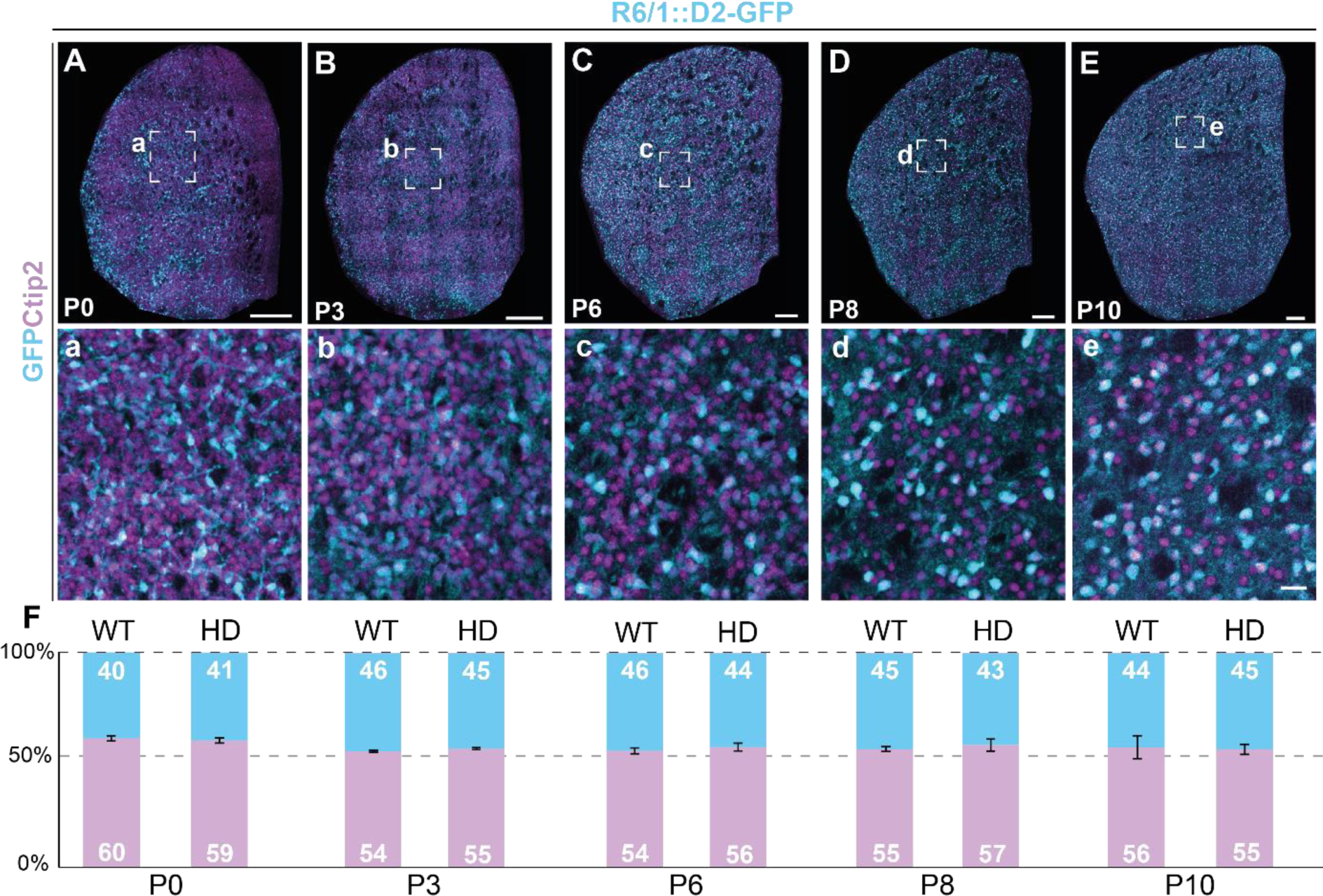
Distribution of GFP^+^ neurons in the striatum of WT::- and R6/1::D2-GFP mice during the first two postnatal weeks. **A-E.** Representative images of GFP and Ctip2 labeling at 5 developmental stages and enlarged images in a-e. F. Quantification of the percentage of GFP^+^ D2-MSNs over the total number of Ctip2+ MSNs for WT and R6/1 mice at 5 developmental stages. Scale bars: A-E, 200 μm; a-e, 25 μm.

**Figure S3:**
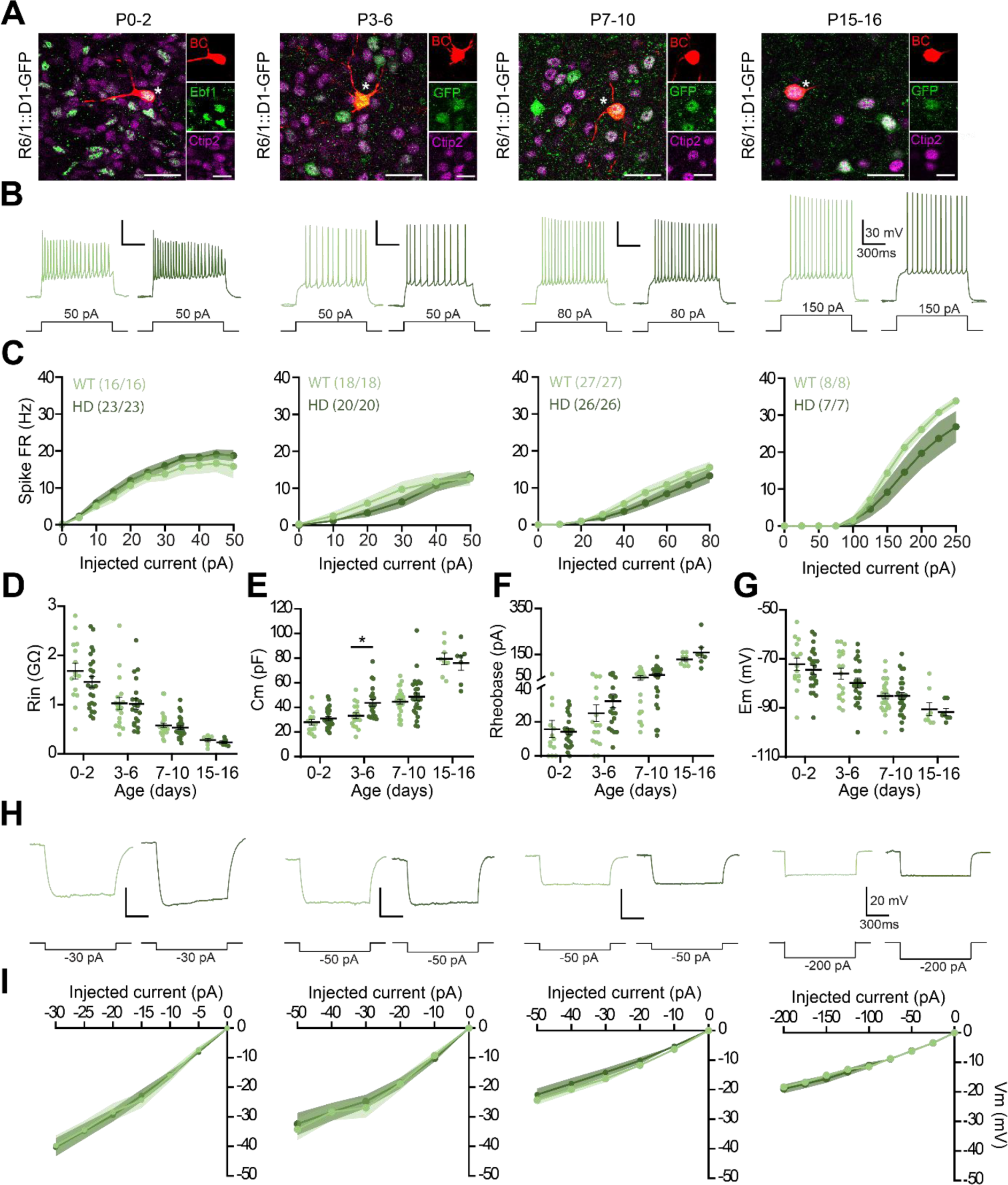
Maturation of the electrophysiological properties of D1-MSNs is not impaired in R6/1 mice during the first two postnatal weeks. **A**. Examples of recorded biocytin-filled neurons at 4 postnatal stages: P0-2, P3-6, P7-10 and P15-16. During the first postnatal days, immunohistochemistry directed against the D1-MSN marker Ebf1 confirmed the D1-MSN nature of the recorded neurons. Recorded neurons are identified by an asterisk. **B**. Representative responses of D1-MSNs in WT mice (light green) or R6/1 mice (dark green) following injection of a depolarizing current. **C**. Mean discharge frequency/injected current curves obtained for each age. **D-G**. Main electrophysiological properties of D1-MSNs with membrane resistance (Rm, **D**), membrane capacitance (Cm, **E**), rheobase (**F**) and resting membrane potential (Em, **G**) plotted longitudinally. **H**. Representative responses of D1-MSNs in WT mice (light green) or R6/1 mice (dark green) following injection of a hyperpolarizing current. **I**. Potential/current curves obtained for each age. Scale bars: 25 and 10 μm.

**Figure S4:**
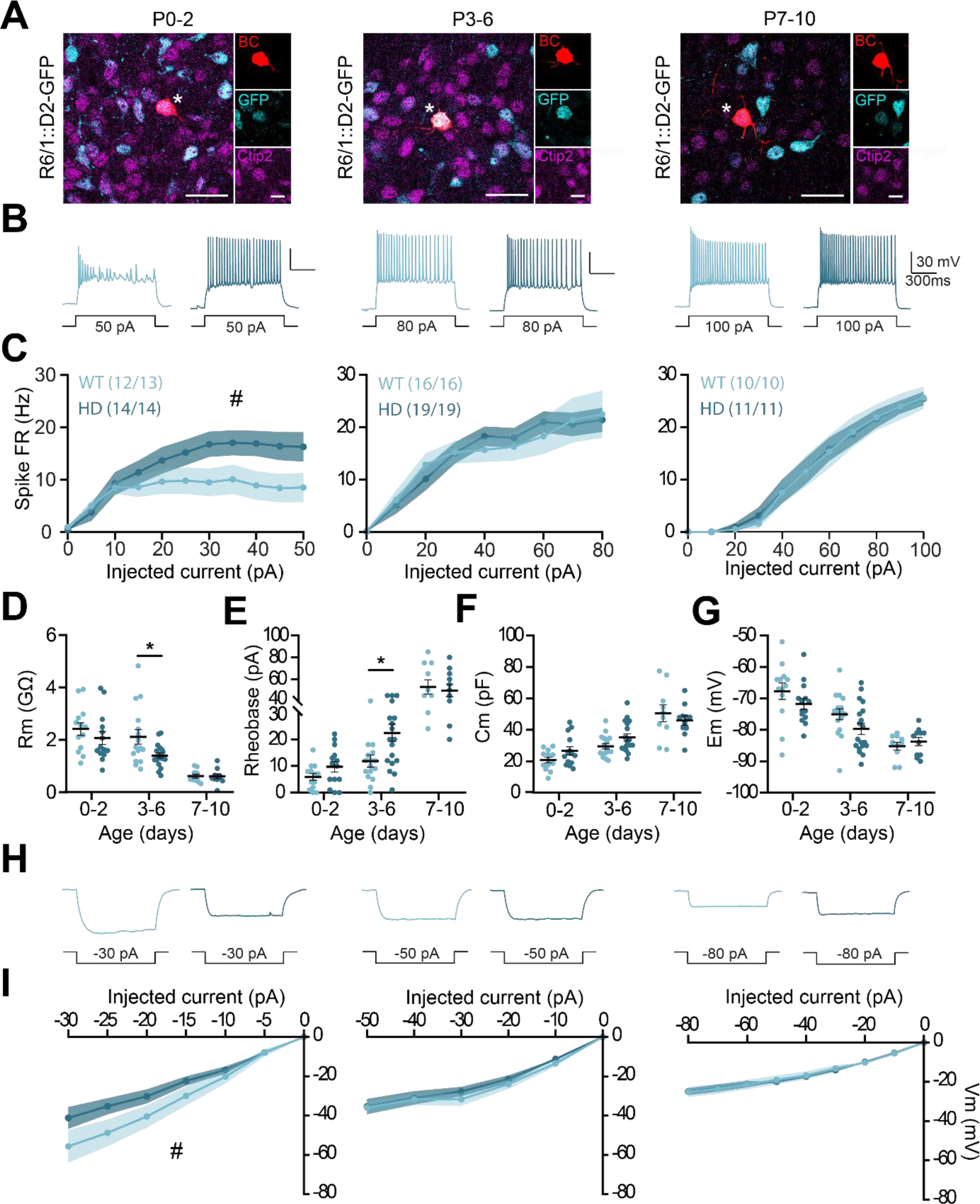
Maturation of the electrophysiological properties of D2-MSNs is impaired in CAG140 mice during the first postnatal days. **A**. Examples of recorded biocytin-filled neurons at 3 postnatal stages: P0-2, P3-6 and P7-10. Recorded neurons are identified by an asterisk. **B**. Representative responses of D2-MSNs in WT mice (light blue) or CAG140 mice (dark blue) following injection of a depolarizing current. **C**. Mean discharge frequency/injected current curves obtained for each age. **D-G**. Main electrophysiological properties of D2-MSNs with membrane input resistance (Rin, **D**), rheobase (**E**), capacitance (Cm, F) and resting membrane potential (Em, **G**) plotted longitudinally. **H**. Representative responses of D2-MSNs in WT mice (light blue) or CAG140 mice (dark blue) following injection of a hyperpolarizing current. **I**. Potential/current curves obtained for each age. Scale bars: 25 and 10 μm.

**Figure S5:**
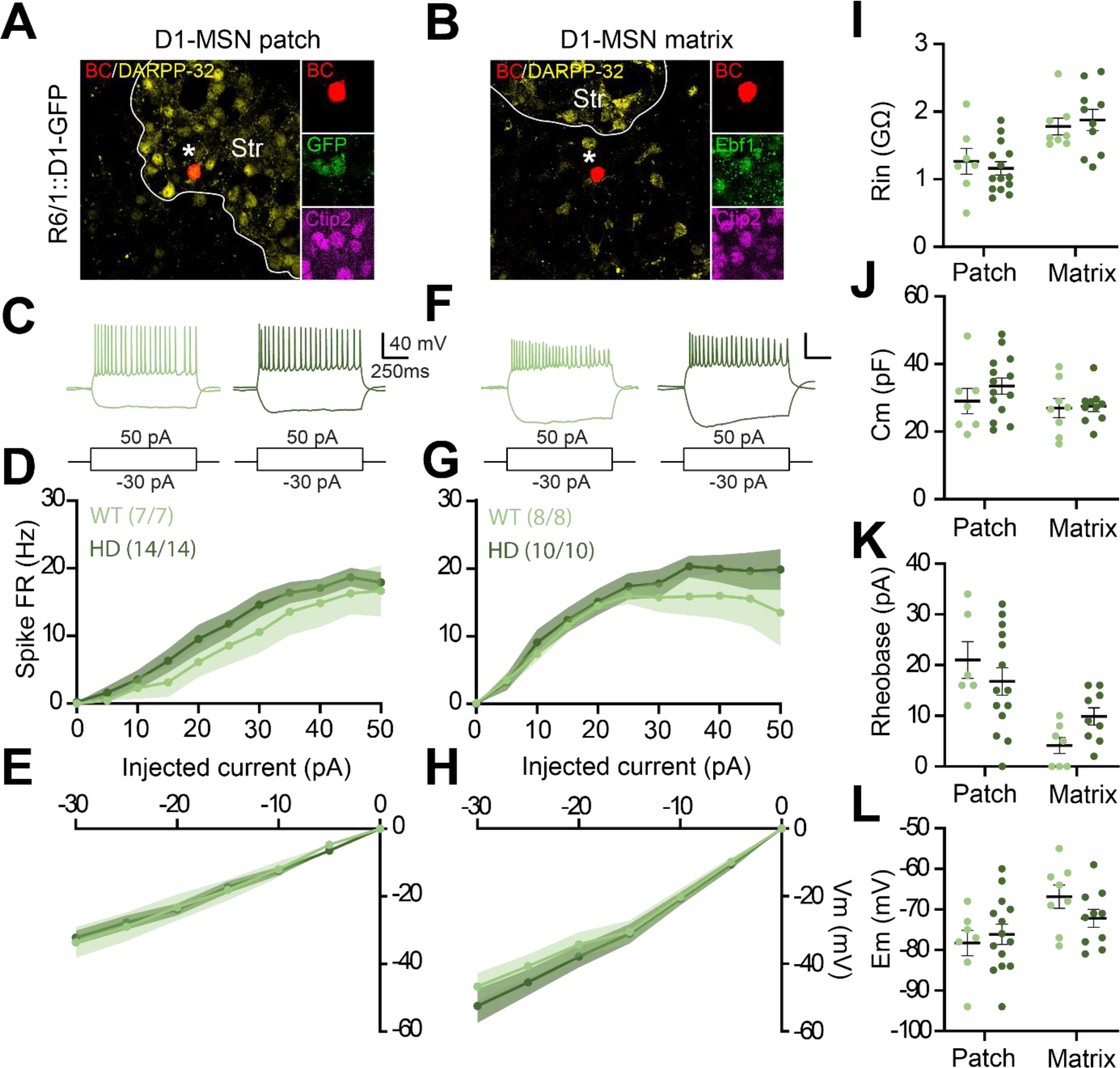
Electrophysiological properties of D1-MSNs from R6/1 mice are not altered in a compartment-dependent manner during the first postnatal days. A-B. Examples of D1-MSNs neurons in either striosome (**A**; GFP+/DARPP32^+^) matrix (**B**; Ebf1+/DARPP32-). Recorded neurons are identified by an asterisk. **C**, **F**. Representative D1-MSNs responses from striosome (**C**) or matrix (**F**) in WT (light green) or R6/1 (dark green) mice following the injection of hyperpolarizing and depolarizing currents. **D, G**. Mean discharge frequency/injected current curves obtained for each condition. **E, H**. Membrane potential/ current curves obtained for each condition. **I-L**. Main electrophysiological parameters of D1-MSNs with resistance membrane resistance (Em, **I**), membrane capacitance (Cm, **J**), rheobase (**K**) and resting membrane potential (Em, **L**) as a function of the striatal compartment. Scale bars: 25 and 10 μm.

**Figure S6:**
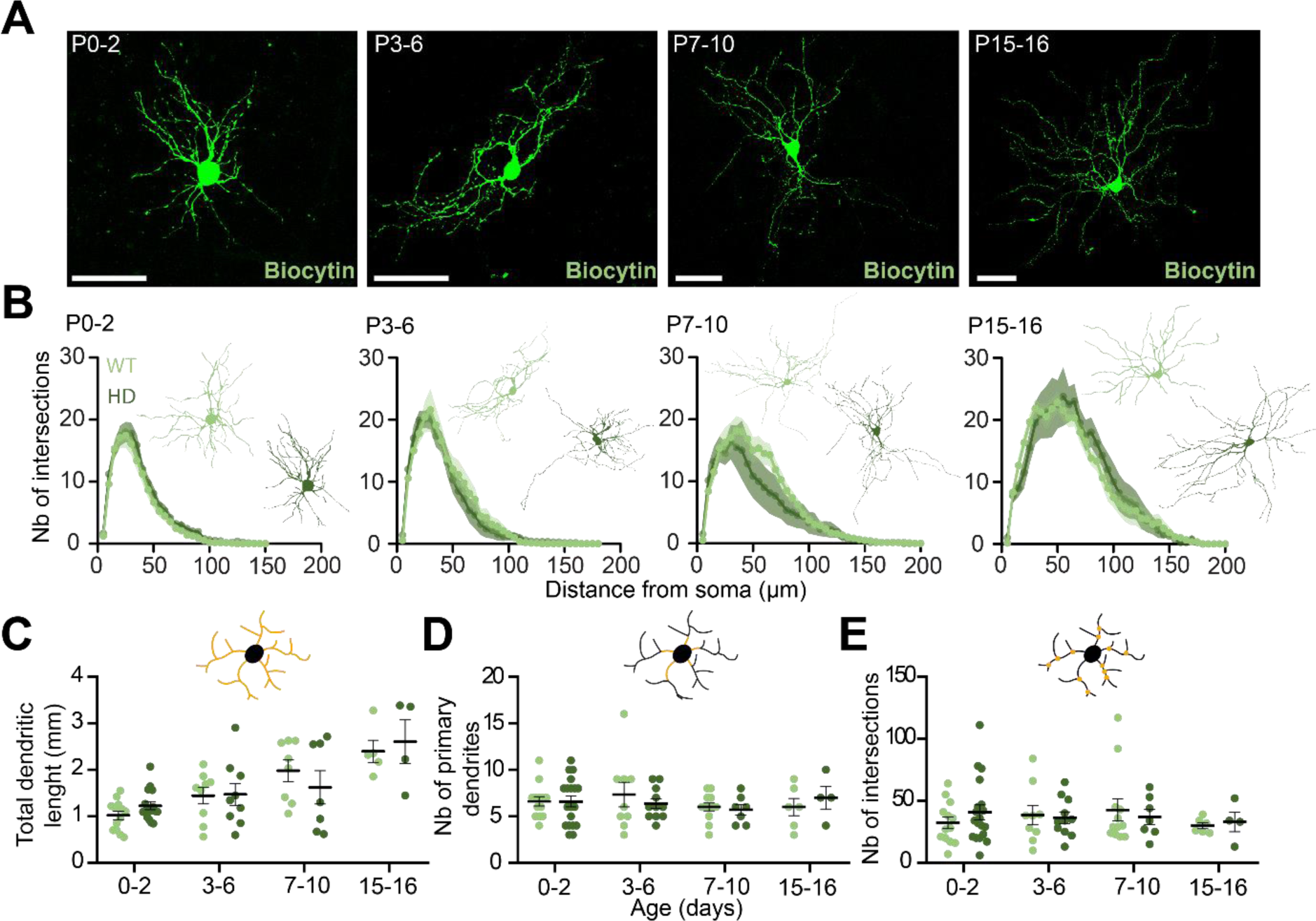
Maturation of the morphological properties of D1-MSNs is not impaired in R6/1 mice during the first two postnatal weeks. **A.** Examples of biocytin-filled D1-MSNs at different postnatal stages. **B.** Scholl analysis illustrating the number of intersections between the dendritic tree and concentric circles spaced 5 μm from the soma. **C-E.** Main features of the dendritic arborization with dendritic length **(C)**, number of primary dendrites **(D)** and number of nodes **(E)**.

**Figure S7:**
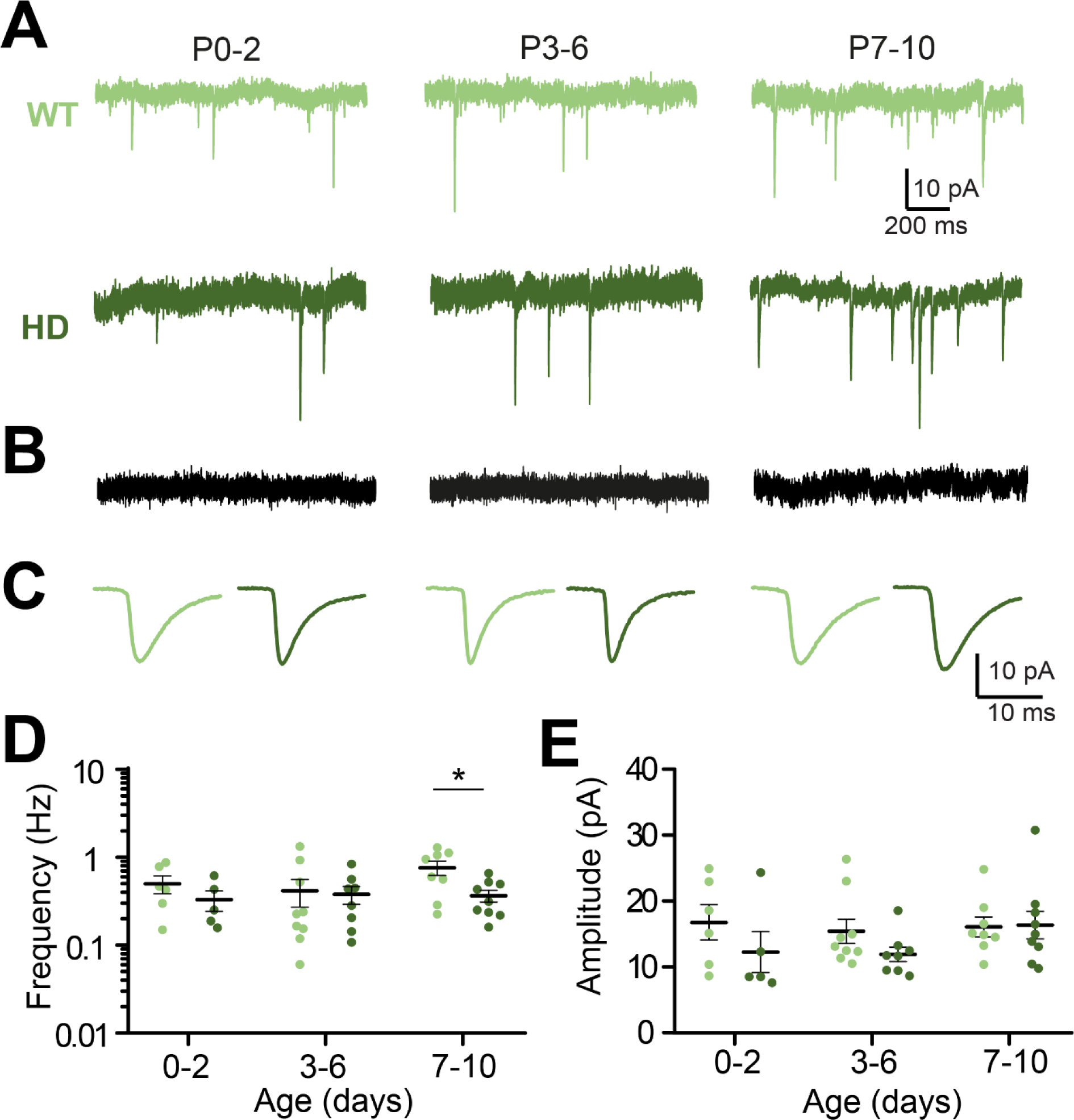
Glutamatergic transmission is impaired in D1-MSNs from R6/1 mice during the first two postnatal weeks. **A.** Representative recordings of spontaneous excitatory postsynaptic currents (sEPSC) imposed at a potential of -70 mV in WT (top, light green) and R6/1 (bottom, dark green) mice at P0-2, P3-6 and P7-10, recorded in the presence of GABA_A_ and GABA_B_ antagonists (5 µM GABAzine + 1µM CGP55485). **B**. sEPSCs disappear when glutamatergic antagonists (20 µM DNQX + 50 µM APV) are added to the bath. **C**. Average traces of sEPSCs obtained for each condition. **D-E**. Mean frequency (**D**) and amplitude (**E**) of sEPSCs as a function of age.

**Figure S8:**
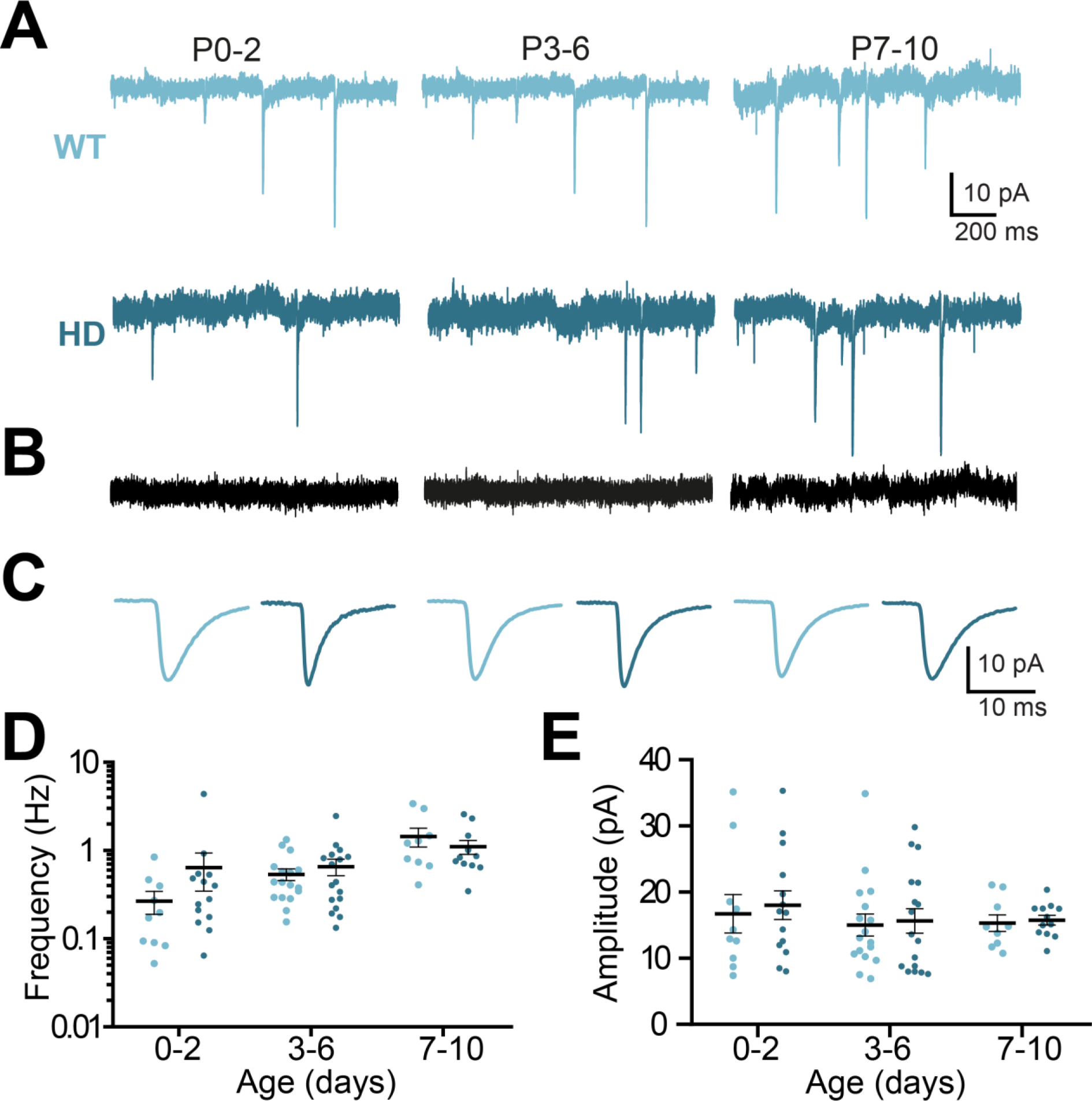
Glutamatergic transmission is not impaired in D2-MSNs from CAG140 mice during the first two postnatal weeks. **A.** Representative recordings of spontaneous excitatory postsynaptic currents (sEPSC) imposed at a potential of -70 mV in WT (top, light blue) and CAG140 (bottom, dark blue) mice at P0-2, P3-6 and P7-10, recorded in the presence of GABA_A_ and GABA_B_ antagonists (5 µM GABAzine + 1µM CGP55485). **B**. sEPSCs disappear when glutamatergic antagonists (20 µM DNQX + 50 µM APV) are added to the bath. **C**. Average traces of sEPSCs obtained for each condition. **D-E**. Mean frequency (**D**) and amplitude (**E**) of sEPSCs as a function of age.

**Figure S9:**
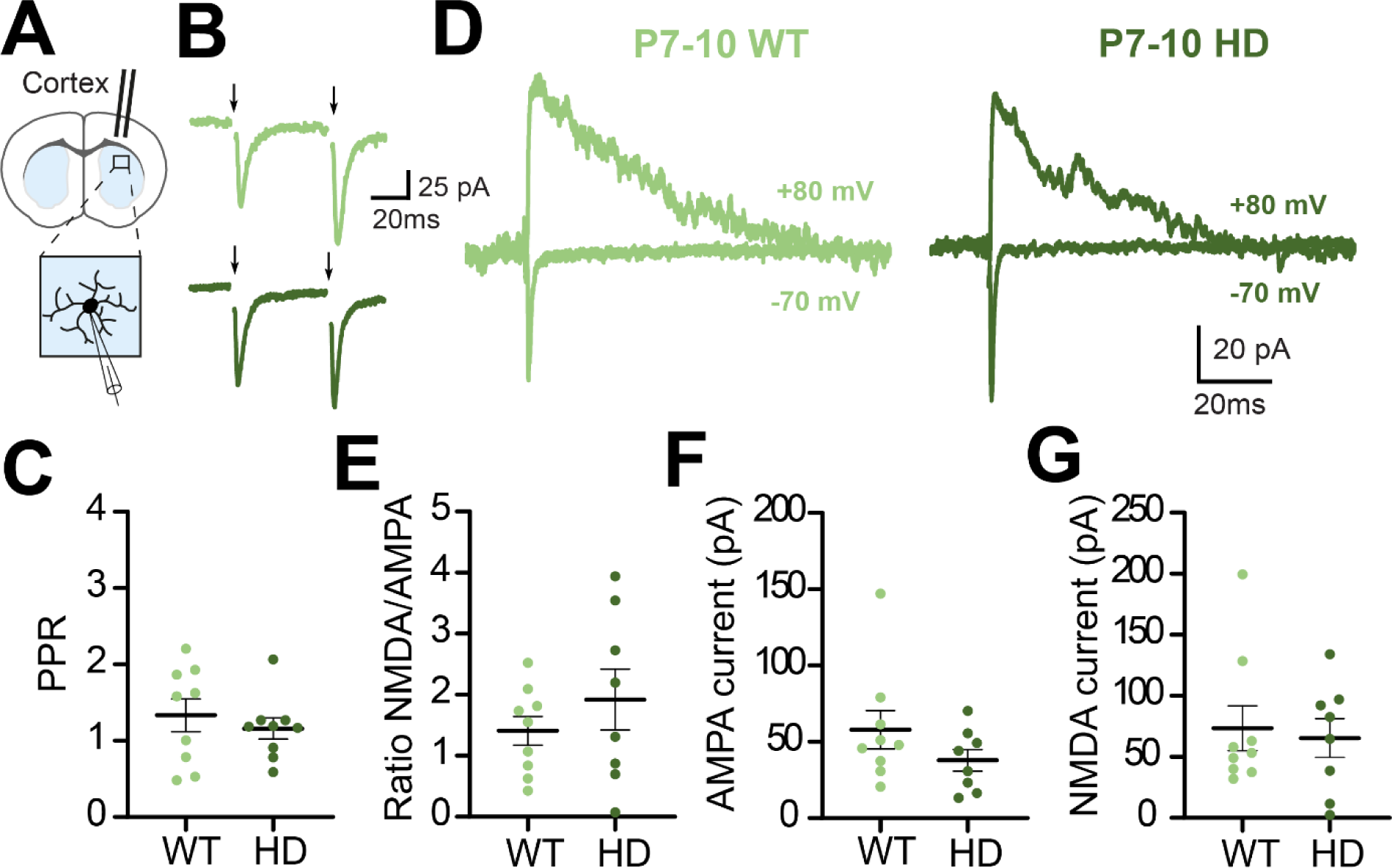
Cortico-striatal transmission on D1-MSNs is not impaired in R6/1 mice during the second postnatal week. **A.** Schematic of the experiment showing the position of the stimulating electrode and the recording electrode. **B.** Representative recordings of evoked excitatory postsynaptic currents (eEPSCs) at a potential of -70 mV in WT (light green) and R6/1 (dark green) mice. **C.** Average paired pulse ratio (EPSC2/EPSC1). **D.** Representative recordings at an imposed potential of either -70 mV to record AMPA receptor-mediated currents or at +80 mV to record currents mediated by both AMPA and NMDA receptors. **E.** NMDA/AMPA ratio obtained by dividing the amplitude values of NMDA currents by those of AMPA currents. **F-G.** Amplitude values of AMPA (F) and NMDA (G) currents. All recordings were performed in presence of GABAA and GABAB receptors antagonists (5 µM GABAzine + 1 µM CGP55845).

## References

Ahmed, N. Y., Knowles, R., Liu, L., Yan, Y., Li, X., Schumann, U., Wang, Y., Sontani, Y., Reynolds, N., Natoli, R., Wen, J., Del Pino, I., Mi, D., & Dehorter, N. (2023). Developmental deficits of MGE-derived interneurons in the Cntnap2 knockout mouse model of autism spectrum disorder. Frontiers in Cell and Developmental Biology, 11, 1112062. 10.3389/fcell.2023.1112062

Albin, R. L., Reiner, A., Anderson, K. D., Dure, L. S., Handelin, B., Balfour, R., Whetsell, W. O., Penney, J. B., & Young, A. B. (1992). Preferential loss of striato-external pallidal projection neurons in presymptomatic Huntington’s disease. Annals of Neurology, 31(4), 425–430. 10.1002/ana.410310412

Araki, K. Y., Sims, J. R., & Bhide, P. G. (2007). Dopamine receptor mRNA and protein expression in the mouse corpus striatum and cerebral cortex during pre- and postnatal development. Brain Research, 1156, 31–45. 10.1016/j.brainres.2007.04.043

Arlotta, P., Molyneaux, B. J., Jabaudon, D., Yoshida, Y., & Macklis, J. D. (2008). Ctip2 controls the differentiation of medium spiny neurons and the establishment of the cellular architecture of the striatum. The Journal of Neuroscience : The Official Journal of the Society for Neuroscience, 28(3), 622–632. 10.1523/JNEUROSCI.2986-07.2008

Arteaga-Bracho, E. E., Gulinello, M., Winchester, M. L., Pichamoorthy, N., Petronglo, J. R., Zambrano, A. D., Inocencio, J., De Jesus, C. D., Louie, J. O., Gokhan, S., Mehler, M. F., & Molero, A. E. (2016). Postnatal and adult consequences of loss of huntingtin during development: Implications for Huntington’s disease. Neurobiology of Disease, 96, 144–155. 10.1016/j.nbd.2016.09.006

Bankhead, P., Loughrey, M. B., Fernández, J. A., Dombrowski, Y., McArt, D. G., Dunne, P. D., McQuaid, S., Gray, R. T., Murray, L. J., Coleman, H. G., James, J. A., Salto-Tellez, M., & Hamilton, P. W. (2017). QuPath: Open source software for digital pathology image analysis. Scientific Reports, 7(1), 16878. 10.1038/s41598-017-17204-5

Barnat, M., Capizzi, M., Aparicio, E., Boluda, S., Wennagel, D., Kacher, R., Kassem, R., Lenoir, S., Agasse, F., Braz, B. Y., Liu, J.-P., Ighil, J., Tessier, A., Zeitlin, S. O., Duyckaerts, C., Dommergues, M., Durr, A., & Humbert, S. (2020). Huntington’s disease alters human neurodevelopment. *Science (New York*, N.Y.), 369(6505), 787–793. 10.1126/science.aax3338

Barnat, M., Le Friec, J., Benstaali, C., & Humbert, S. (2017). Huntingtin-Mediated Multipolar-Bipolar Transition of Newborn Cortical Neurons Is Critical for Their Postnatal Neuronal Morphology. Neuron, 93(1), 99–114. 10.1016/j.neuron.2016.11.035

Baydyuk, M., Russell, T., Liao, G.-Y., Zang, K., An, J. J., Reichardt, L. F., & Xu, B. (2011). TrkB receptor controls striatal formation by regulating the number of newborn striatal neurons. Proceedings of the National Academy of Sciences of the United States of America, 108(4), 1669–1674. 10.1073/pnas.1004744108

Baydyuk, M., & Xu, B. (2014). BDNF signaling and survival of striatal neurons. Frontiers in Cellular Neuroscience, 8, 254. 10.3389/fncel.2014.00254

Bitzenhofer, S. H., Pöpplau, J. A., Chini, M., Marquardt, A., & Hanganu-Opatz, I. L. (2021). A transient developmental increase in prefrontal activity alters network maturation and causes cognitive dysfunction in adult mice. Neuron, 109(8), 1350–1364.e6. 10.1016/j.neuron.2021.02.011

Blumenstock, S., & Dudanova, I. (2020). Cortical and Striatal Circuits in Huntington’s Disease. Frontiers in Neuroscience, 14, 82. 10.3389/fnins.2020.00082

Braz, B. Y., Wennagel, D., Ratié, L., de Souza, D. A. R., Deloulme, J. C., Barbier, E. L., Buisson, A., Lanté, F., & Humbert, S. (2022). Treating early postnatal circuit defect delays Huntington’s disease onset and pathology in mice. *Science (New York*, N.Y.), 377(6613), eabq5011. 10.1126/science.abq5011

Caille, I., Dumartin, B., Le Moine, C., Begueret, J., & Bloch, B. (1995). Ontogeny of the D1 dopamine receptor in the rat striatonigral system: an immunohistochemical study. The European Journal of Neuroscience, 7(4), 714– 722. 10.1111/j.1460-9568.1995.tb00675.x

Capizzi, M., Carpentier, R., Denarier, E., Adrait, A., Kassem, R., Mapelli, M., Couté, Y., & Humbert, S. (2022). Developmental defects in Huntington’s disease show that axonal growth and microtubule reorganization require NUMA1. Neuron, 110(1), 36–50.e5. 10.1016/j.neuron.2021.10.033

Carrillo-Reid, L., Day, M., Xie, Z., Melendez, A. E., Kondapalli, J., Plotkin, J. L., Wokosin, D. L., Chen, Y., Kress, G. J., Kaplitt, M., Ilijic, E., Guzman, J. N., Chan, C. S., & Surmeier, D. J. (2019). Mutant huntingtin enhances activation of dendritic Kv4 K+ channels in striatal spiny projection neurons. ELife, 8. 10.7554/eLife.40818

Crandall, J. E., McCarthy, D. M., Araki, K. Y., Sims, J. R., Ren, J.-Q., & Bhide, P. G. (2007). Dopamine receptor activation modulates GABA neuron migration from the basal forebrain to the cerebral cortex. The Journal of Neuroscience : The Official Journal of the Society for Neuroscience, 27(14), 3813–3822. 10.1523/JNEUROSCI.5124-06.2007

Crittenden, J. R., & Graybiel, A. M. (2011). Basal Ganglia disorders associated with imbalances in the striatal striosome and matrix compartments. Frontiers in Neuroanatomy, 5, 59. 10.3389/fnana.2011.00059

Dehorter, N., & Del Pino, I. (2020). Shifting Developmental Trajectories During Critical Periods of Brain Formation. Frontiers in Cellular Neuroscience, 14, 283. 10.3389/fncel.2020.00283

Dehorter, N., Michel, F. J., Marissal, T., Rotrou, Y., Matrot, B., Lopez, C., Humphries, M. D., & Hammond, C. (2011). Onset of Pup Locomotion Coincides with Loss of NR2C/D-Mediated Cortico-Striatal EPSCs and Dampening of Striatal Network Immature Activity. Frontiers in Cellular Neuroscience, 5, 24. 10.3389/fncel.2011.00024

Deng, Y. P., Albin, R. L., Penney, J. B., Young, A. B., Anderson, K. D., & Reiner, A. (2004). Differential loss of striatal projection systems in Huntington’s disease: a quantitative immunohistochemical study. Journal of Chemical Neuroanatomy, 27(3), 143–164. 10.1016/j.jchemneu.2004.02.005

Duyao, M. P., Auerbach, A. B., Ryan, A., Persichetti, F., Barnes, G. T., McNeil, S. M., Ge, P., Vonsattel, J. P., Gusella, J. F., & Joyner, A. L. (1995). Inactivation of the mouse Huntington’s disease gene homolog Hdh. *Science (New York*, N.Y.), 269(5222), 407–410. 10.1126/science.7618107

Fishell, G., & van der Kooy, D. (1989). Pattern formation in the striatum: developmental changes in the distribution of striatonigral projections. Brain Research. Developmental Brain Research, 45(2), 239–255. 10.1016/0165-3806(89)90042-4

Fishell, G., & van der Kooy, D. (1991). Pattern formation in the striatum: neurons with early projections to the substantia nigra survive the cell death period. The Journal of Comparative Neurology, 312(1), 33–42. 10.1002/cne.903120104

Flaherty, A. W., & Graybiel, A. M. (1994). Input-output organization of the sensorimotor striatum in the squirrel monkey. The Journal of Neuroscience : The Official Journal of the Society for Neuroscience, 14(2), 599–610. 10.1523/JNEUROSCI.14-02-00599.1994

Friedman, A., Homma, D., Gibb, L. G., Amemori, K.-I., Rubin, S. J., Hood, A. S., Riad, M. H., & Graybiel, A. M. (2015). A Corticostriatal Path Targeting Striosomes Controls Decision-Making under Conflict. Cell, 161(6), 1320–1333. 10.1016/j.cell.2015.04.049

Gagnon, D., Petryszyn, S., Sanchez, M. G., Bories, C., Beaulieu, J. M., De Koninck, Y., Parent, A., & Parent, M. (2017). Striatal Neurons Expressing D1 and D2 Receptors are Morphologically Distinct and Differently Affected by Dopamine Denervation in Mice. Scientific Reports, 7, 41432. 10.1038/srep41432

Garel, S., Marín, F., Mattéi, M. G., Vesque, C., Vincent, A., & Charnay, P. (1997). Family of Ebf/Olf-1-related genes potentially involved in neuronal differentiation and regional specification in the central nervous system. Developmental Dynamics : An Official Publication of the American Association of Anatomists, 210(3), 191–205. 10.1002/(SICI)1097-0177(199711)210:3<191::AID-AJA1>3.0.CO;2-B

Ginés, S., Bosch, M., Marco, S., Gavaldà, N., Díaz-Hernández, M., Lucas, J. J., Canals, J. M., & Alberch, J. (2006). Reduced expression of the TrkB receptor in Huntington’s disease mouse models and in human brain. The European Journal of Neuroscience, 23(3), 649–658. 10.1111/j.1460-9568.2006.04590.x

He, Q., Arroyo, E. D., Smukowski, S. N., Xu, J., Piochon, C., Savas, J. N., Portera-Cailliau, C., & Contractor, A. (2019). Critical period inhibition of NKCC1 rectifies synapse plasticity in the somatosensory cortex and restores adult tactile response maps in fragile X mice. Molecular Psychiatry, 24(11), 1732–1747. 10.1038/s41380-018-0048-y

Jung, A. B., & Bennett, J. P. (1996). Development of striatal dopaminergic function. I. Pre- and postnatal development of mRNAs and binding sites for striatal D1 (D1a) and D2 (D2a) receptors. *Brain Research. Developmental Brain Research*, *94*(2), 109–120. 10.1016/0165-3806(96)00033-8

Kirischuk, S., Sinning, A., Blanquie, O., Yang, J.-W., Luhmann, H. J., & Kilb, W. (2017). Modulation of Neocortical Development by Early Neuronal Activity: Physiology and Pathophysiology. Frontiers in Cellular Neuroscience, 11, 379. 10.3389/fncel.2017.00379

Kita, T., Kita, H., & Kitai, S. T. (1984). Passive electrical membrane properties of rat neostriatal neurons in an in vitro slice preparation. Brain Research, 300(1), 129–139. 10.1016/0006-8993(84)91347-7

Kozorovitskiy, Y., Saunders, A., Johnson, C. A., Lowell, B. B., & Sabatini, B. L. (2012). Recurrent network activity drives striatal synaptogenesis. Nature, 485(7400), 646–650. 10.1038/nature11052

Krajeski, R. N., Macey-Dare, A., van Heusden, F., Ebrahimjee, F., & Ellender, T. J. (2019). Dynamic postnatal development of the cellular and circuit properties of striatal D1 and D2 spiny projection neurons. The Journal of Physiology, 597(21), 5265–5293. 10.1113/JP278416

Lebouc, M., Richard, Q., Garret, M., & Baufreton, J. (2020). Striatal circuit development and its alterations in Huntington’s disease. Neurobiology of Disease, 145, 105076. 10.1016/j.nbd.2020.105076

Lee, T., Kaneko, T., Taki, K., & Mizuno, N. (1997). Preprodynorphin-, preproenkephalin-, and preprotachykinin-expressing neurons in the rat neostriatum: an analysis by immunocytochemistry and retrograde tracing. The Journal of Comparative Neurology, 386(2), 229–244. 10.1002/(sici)1096-9861(19970922)386:2<229::aid-cne5>3.0.co;2-3

Mandal, M., Wei, J., Zhong, P., Cheng, J., Duffney, L. J., Liu, W., Yuen, E. Y., Twelvetrees, A. E., Li, S., Li, X.-J., Kittler, J. T., & Yan, Z. (2011). Impaired alpha-amino-3-hydroxy-5-methyl-4-isoxazolepropionic acid (AMPA) receptor trafficking and function by mutant huntingtin. The Journal of Biological Chemistry, 286(39), 33719–33728. 10.1074/jbc.M111.236521

Mangiarini, L., Sathasivam, K., Seller, M., Cozens, B., Harper, A., Hetherington, C., Lawton, M., Trottier, Y., Lehrach, H., Davies, S. W., & Bates, G. P. (1996). Exon 1 of the HD gene with an expanded CAG repeat is sufficient to cause a progressive neurological phenotype in transgenic mice. Cell, 87(3), 493–506. 10.1016/s0092-8674(00)81369-0

Matsushima, A., Pineda, S. S., Crittenden, J. R., Lee, H., Galani, K., Mantero, J., Tombaugh, G., Kellis, M., Heiman, M., & Graybiel, A. M. (2023). Transcriptional vulnerabilities of striatal neurons in human and rodent models of Huntington’s disease. Nature Communications, 14(1), 282. 10.1038/s41467-022-35752-x

McKinstry, S. U., Karadeniz, Y. B., Worthington, A. K., Hayrapetyan, V. Y., Ozlu, M. I., Serafin-Molina, K., Risher, W. C., Ustunkaya, T., Dragatsis, I., Zeitlin, S., Yin, H. H., & Eroglu, C. (2014). Huntingtin is required for normal excitatory synapse development in cortical and striatal circuits. The Journal of Neuroscience : The Official Journal of the Society for Neuroscience, 34(28), 9455–9472. 10.1523/JNEUROSCI.4699-13.2014

Merchan-Sala, P., Nardini, D., Waclaw, R. R., & Campbell, K. (2017). Selective neuronal expression of the SoxE factor, Sox8, in direct pathway striatal projection neurons of the developing mouse brain. *The Journal of Comparative Neurology*, *525*(13), 2805–2819. 10.1002/cne.24232

Molero, A. E., Arteaga-Bracho, E. E., Chen, C. H., Gulinello, M., Winchester, M. L., Pichamoorthy, N., Gokhan, S., Khodakhah, K., & Mehler, M. F. (2016). Selective expression of mutant huntingtin during development recapitulates characteristic features of Huntington’s disease. Proceedings of the National Academy of Sciences of the United States of America, 113(20), 5736–5741. 10.1073/pnas.1603871113

Molina-Calavita, M., Barnat, M., Elias, S., Aparicio, E., Piel, M., & Humbert, S. (2014). Mutant huntingtin affects cortical progenitor cell division and development of the mouse neocortex. The Journal of Neuroscience : The Official Journal of the Society for Neuroscience, 34(30), 10034–10040. 10.1523/JNEUROSCI.0715-14.2014

Morton, A. J., Nicholson, L. F., & Faull, R. L. (1993). Compartmental loss of NADPH diaphorase in the neuropil of the human striatum in Huntington’s disease. Neuroscience, 53(1), 159–168. 10.1016/0306-4522(93)90294-p

Morton, A. J., Skillings, E. A., Wood, N. I., & Zheng, Z. (2019). Antagonistic pleiotropy in mice carrying a CAG repeat expansion in the range causing Huntington’s disease. Scientific Reports, 9(1), 37. 10.1038/s41598-018-37102-8

Nasir, J., Floresco, S. B., O’Kusky, J. R., Diewert, V. M., Richman, J. M., Zeisler, J., Borowski, A., Marth, J. D., Phillips, A. G., & Hayden, M. R. (1995). Targeted disruption of the Huntington’s disease gene results in embryonic lethality and behavioral and morphological changes in heterozygotes. Cell, 81(5), 811–823. 10.1016/0092-8674(95)90542-1

Nisenbaum, E. S., Wilson, C. J., Foehring, R. C., & Surmeier, D. J. (1996). Isolation and characterization of a persistent potassium current in neostriatal neurons. Journal of Neurophysiology, 76(2), 1180–1194. 10.1152/jn.1996.76.2.1180

Peixoto, R. T., Wang, W., Croney, D. M., Kozorovitskiy, Y., & Sabatini, B. L. (2016). Early hyperactivity and precocious maturation of corticostriatal circuits in Shank3B(-/-) mice. Nature Neuroscience, 19(5), 716–724. 10.1038/nn.4260

Pert, C. B., Kuhar, M. J., & Snyder, S. H. (1976). Opiate receptor: autoradiographic localization in rat brain. Proceedings of the National Academy of Sciences of the United States of America, 73(10), 3729–3733. 10.1073/pnas.73.10.3729

Pouladi, M. A., Morton, A. J., & Hayden, M. R. (2013). Choosing an animal model for the study of Huntington’s disease. Nature Reviews. Neuroscience, 14(10), 708– 721. 10.1038/nrn3570

Rao, P. A., Molinoff, P. B., & Joyce, J. N. (1991). Ontogeny of dopamine D1 and D2 receptor subtypes in rat basal ganglia: a quantitative autoradiographic study. Brain Research. Developmental Brain Research, 60(2), 161–177. 10.1016/0165-3806(91)90045-k

Ratié, L., & Humbert, S. (2024). A developmental component to Huntington’s disease. Revue Neurologique. 10.1016/j.neurol.2024.04.001

Reiner, A., Albin, R. L., Anderson, K. D., D’Amato, C. J., Penney, J. B., & Young, A. B. (1988). Differential loss of striatal projection neurons in Huntington disease. Proceedings of the National Academy of Sciences of the United States of America, 85(15), 5733–5737. 10.1073/pnas.85.15.5733

Rogalski, S. L., Appleyard, S. M., Pattillo, A., Terman, G. W., & Chavkin, C. (2000). TrkB activation by brain-derived neurotrophic factor inhibits the G protein-gated inward rectifier Kir3 by tyrosine phosphorylation of the channel. The Journal of Biological Chemistry, 275(33), 25082–25088. 10.1074/jbc.M000183200

Ross, C. A., Reilmann, R., Cardoso, F., McCusker, E. A., Testa, C. M., Stout, J. C., Leavitt, B. R., Pei, Z., Landwehrmeyer, B., Martinez, A., Levey, J., Srajer, T., Bang, J., & Tabrizi, S. J. (2019). Movement Disorder Society Task Force Viewpoint: Huntington’s Disease Diagnostic Categories. Movement Disorders Clinical Practice, 6(7), 541–546. 10.1002/mdc3.12808

Saudou, F., & Humbert, S. (2016). The Biology of Huntingtin. Neuron, 89(5), 910– 926. 10.1016/j.neuron.2016.02.003

Sharott, A., Vinciati, F., Nakamura, K. C., & Magill, P. J. (2017). A Population of Indirect Pathway Striatal Projection Neurons Is Selectively Entrained to Parkinsonian Beta Oscillations. The Journal of Neuroscience : The Official Journal of the Society for Neuroscience, 37(41), 9977–9998. 10.1523/JNEUROSCI.0658-17.2017

Smith, J. B., Klug, J. R., Ross, D. L., Howard, C. D., Hollon, N. G., Ko, V. I., Hoffman, H., Callaway, E. M., Gerfen, C. R., & Jin, X. (2016). Genetic-Based Dissection Unveils the Inputs and Outputs of Striatal Patch and Matrix Compartments. Neuron, 91(5), 1069–1084. 10.1016/j.neuron.2016.07.046

Smith, Y., Raju, D. V, Pare, J.-F., & Sidibe, M. (2004). The thalamostriatal system: a highly specific network of the basal ganglia circuitry. Trends in Neurosciences, 27(9), 520–527. 10.1016/j.tins.2004.07.004

Tabrizi, S. J., Schobel, S., Gantman, E. C., Mansbach, A., Borowsky, B., Konstantinova, P., Mestre, T. A., Panagoulias, J., Ross, C. A., Zauderer, M., Mullin, A. P., Romero, K., Sivakumaran, S., Turner, E. C., Long, J. D., Sampaio, C., & Huntington’s Disease Regulatory Science Consortium (HD-RSC). (2022). A biological classification of Huntington’s disease: the Integrated Staging System. The Lancet. Neurology, 21(7), 632–644. 10.1016/S1474-4422(22)00120-X

Tepper, J. M., & Trent, F. (1993). In vivo studies of the postnatal development of rat neostriatal neurons. Progress in Brain Research, 99, 35–50. 10.1016/s0079-6123(08)61337-0

Tereshchenko, A., van der Plas, E., Mathews, K. D., Epping, E., Conrad, A. L., Langbehn, D. R., & Nopoulos, P. (2020). Developmental Trajectory of Height, Weight, and BMI in Children and Adolescents at Risk for Huntington’s Disease: Effect of mHTT on Growth. Journal of Huntington’s Disease, 9(3), 245–251. 10.3233/JHD-200407

Tinterri, A., Menardy, F., Diana, M. A., Lokmane, L., Keita, M., Coulpier, F., Lemoine, S., Mailhes, C., Mathieu, B., Merchan-Sala, P., Campbell, K., Gyory, I., Grosschedl, R., Popa, D., & Garel, S. (2018). Active intermixing of indirect and direct neurons builds the striatal mosaic. Nature Communications, 9(1), 4725. 10.1038/s41467-018-07171-4

van der Plas, E., Langbehn, D. R., Conrad, A. L., Koscik, T. R., Tereshchenko, A., Epping, E. A., Magnotta, V. A., & Nopoulos, P. C. (2019). Abnormal brain development in child and adolescent carriers of mutant huntingtin. Neurology, 93(10), e1021–e1030. 10.1212/WNL.0000000000008066

Vonsattel, J. P., Myers, R. H., Stevens, T. J., Ferrante, R. J., Bird, E. D., & Richardson, E. P. (1985). Neuropathological classification of Huntington’s disease. Journal of Neuropathology and Experimental Neurology, 44(6), 559–577. 10.1097/00005072-198511000-00003

Voorn, P., Vanderschuren, L. J. M. J., Groenewegen, H. J., Robbins, T. W., & Pennartz, C. M. A. (2004). Putting a spin on the dorsal-ventral divide of the striatum. Trends in Neurosciences, 27(8), 468–474. 10.1016/j.tins.2004.06.006

Yang, L., Su, Z., Wang, Z., Li, Z., Shang, Z., Du, H., Liu, G., Qi, D., Yang, Z., Xu, Z., & Zhang, Z. (2021). Transcriptional profiling reveals the transcription factor networks regulating the survival of striatal neurons. Cell Death & Disease, 12(3), 262. 10.1038/s41419-021-03552-8

Zeitlin, S., Liu, J. P., Chapman, D. L., Papaioannou, V. E., & Efstratiadis, A. (1995). Increased apoptosis and early embryonic lethality in mice nullizygous for the Huntington’s disease gene homologue. Nature Genetics, 11(2), 155–163. 10.1038/ng1095-155

Zhang, H., Zhang, C., Vincent, J., Zala, D., Benstaali, C., Sainlos, M., Grillo-Bosch, D., Daburon, S., Coussen, F., Cho, Y., David, D. J., Saudou, F., Humeau, Y., & Choquet, D. (2018). Modulation of AMPA receptor surface diffusion restores hippocampal plasticity and memory in Huntington’s disease models. Nature Communications, 9(1), 4272. 10.1038/s41467-018-06675-3

